# DANCE 2.0: Transforming single-cell analysis from black box to transparent workflow

**DOI:** 10.1101/2025.07.17.665427

**Authors:** Jiayuan Ding, Zhongyu Xing, Yixin Wang, Renming Liu, Sheng Liu, Zhi Huang, Wenzhuo Tang, Yuying Xie, James Zou, Xiaojie Qiu, Jian Ma, Guoxian Yu, Jiliang Tang

**Author notes:** **Corresponding authors:** *J.D.* (*), G.Y.* (*), J.M.* (*), J.T. These authors contributed equally to this work (Jiayuan Ding, Zhongyu Xing, Yixin Wang).

## Abstract

Preprocessing is a critical step in single-cell data analysis, yet current practices remain largely a black-box, trial-and-error process driven by user intuition, legacy defaults, and ad hoc heuristics. The optimal combination of steps such as normalization, gene selection, and dimensionality reduction varies across tasks, model architectures, and dataset characteristics, hindering reproducibility and method development. We present DANCE 2.0, an automated and interpretable preprocessing platform featuring two key modules: the Method-Aware Preprocessing (MAP) module, which discovers optimal pipelines for task-specific methods via hierarchical search, and the Dataset-Aware Preprocessing (DAP) module, which recommends pipelines for new datasets via similarity-based matching to a reference atlas. Together, MAP and DAP execute over 325,000 pipeline searches across six major tasks – clustering, cell type annotation, imputation, joint embedding, spatial domain identification, and cell type deconvolution – yielding robust and generalizable recommendations. MAP-recommended pipelines consistently outperform original method defaults, with substantial gains across all tasks. Beyond automation, DANCE 2.0 reveals interpretable preprocessing patterns across tasks, methods, and datasets, transforming preprocessing into a transparent, data-driven process. All resources are openly available at https://github.com/OmicsML/dance to support broad community adoption and future methodological advances.

## Introduction

Single-cell technologies – including single-cell RNA sequencing (scRNA-seq) ^1^, spatial transcriptomics ^2–4^, and multi-omics assays ^5–8^ – provide unprecedented resolution to uncover the heterogeneity of cellular states and interactions. Despite their transformative potential, single-cell datasets are inherently noisy, sparse, high-dimensional, and heterogeneous due to both technical limitations and biological complexity. Noise and sparsity often arise from technical dropouts ^9^ and low RNA capture efficiency ^10^, leading to missing or unreliable gene expression values. High dimensionality results from profiling a large number of genes per cell, creating complex feature spaces that complicate modeling. Biological heterogeneity adds further complexity, reflecting variation in cell type, developmental stage, functional state, and microenvironment. Given these challenges, preprocessing is a critical first step in single-cell analysis. Effective preprocessing enhances signal quality, mitigates noise, and corrects for both technical and biological biases, directly shaping the performance and interpretability of downstream methods across tasks such as clustering ^11^, cell type annotation ^12^, imputation ^13^, joint embedding ^14–16^, spatial pattern identification ^17^, and cell type deconvolution ^2^. These steps determine which biological signals are retained, amplified, or lost before any modeling begins, effectively acting as gatekeepers for downstream discovery. Conversely, suboptimal or inconsistent preprocessing can obscure true biological variation, introduce artifacts, and lead to misleading interpretations. Yet despite its central importance, the design and selection of preprocessing workflows remain a persistent challenge in practice.

Although widely used tools – such as Seurat ^18^, Scater ^19^, and Scanpy ^20^ – provide flexible interfaces for individual preprocessing steps (e.g., normalization ^21^, gene selection ^22^, dimensionality reduction ^23^), they offer little or no guidance for composing tasks- and data-set specific preprocessing pipelines. Analysts are often left to navigate a vast and fragmented landscape of options, relying on trial-and-error experimentation, heuristic intuition, or legacy defaults. Most existing studies ^21,24–27^ evaluate the impact of a single, isolated preprocessing component – such as normalization, gene selection, or dimensionality reduction – on specific tasks like clustering or lineage reconstruction. These analyses are typically narrow in scope and fail to capture how preprocessing choices interact across broader analytical objectives. Although a few studies ^28,29^ attempt broader benchmarking, their coverage of tasks and datasets remain limited, and their findings are often difficult to generalize to new methods or data contexts, limiting both their transparency and practical utility. As a result, our understanding of preprocessing remains fragmented, unsystematic, and difficult to operationalize. Another class of existing work that has recently emerged is biological agents, such as Biomni ^30^, CompBioAgent ^31^, scAgent ^32^. These agents focus on creating an environment that aggregates a wide array of commonly used single-cell tools – for example, different methods for cell type annotation. While useful, these systems do not offer intelligent and context-specific decision-making capabilities. When a user inputs their dataset, they cannot determine the optimal preprocessing pipeline tailored to that specific analysis. Instead, it may recommend standard procedures that are broadly accepted but are not optimal for every dataset or downstream task. Together, these limitations underscore an urgent need for a systematic, transparent, and automated platform that can provide context-aware, interpretable, and reproducible preprocessing guidance tailored to both analytical methods and new datasets.

Here, we introduce DANCE 2.0, which addresses this urgent need by transforming single-cell preprocessing from a trial-and-error process into a systematic, data-driven, and interpretable workflow. It provides actionable, context-specific recommendations tailored to both analytical methods and user datasets, delivered through two core modules: the Method-Aware Preprocessing (MAP) module, which tailors preprocessing to specific downstream methods, and the Dataset-Aware Preprocessing (DAP), which recommends pipelines for new datasets via similarity-based matching. In addition to automation, DANCE 2.0 reveals interpretable patterns in preprocessing choices across tasks, methods, and datasets, advancing preprocessing as a transparent and principled component of single-cell analysis. Altogether, MAP and DAP execute over 325,000 pipeline searches across six major tasks – clustering, cell type annotation, imputation, joint embedding, spatial domain identification, and cell type deconvolution – resulting in a reusable, large-scale resource that will be fully open-sourced to promote transparency and reproducibility in the single-cell community.

## RESULTS

### DANCE 2.0 redefines single-cell preprocessing through transparency and automation

In traditional single-cell analysis workflows, preprocessing is often treated as a black box – assembled from ad hoc combinations with little transparency or understanding of why certain configurations succeed. Manually designed pipelines (e.g., *Preprocessing Pipelines A-E*) yield variable performance, yet the rationale behind their success or failure is typically inaccessible (**Fig. 1a**). DANCE 2.0 addresses this limitation by systematically exploring a structured space of commonly used preprocessing operations and visualizing performance across pipeline combinations (**Fig. 1a**; **Methods**). The resulting landscape reveals consistent patterns: high-performing pipelines frequently include components like *NormalizeTotalLog1p*, whereas low-performing ones often involve *FilterGenesTopK*. These associations are quantitatively summarized through correlation scores, highlighting how DANCE 2.0 opens the “black box” of preprocessing by systematically evaluating pipeline combinations, visualizing their performance, and uncovering interpretable patterns across tasks and datasets.

**Figure 1:**
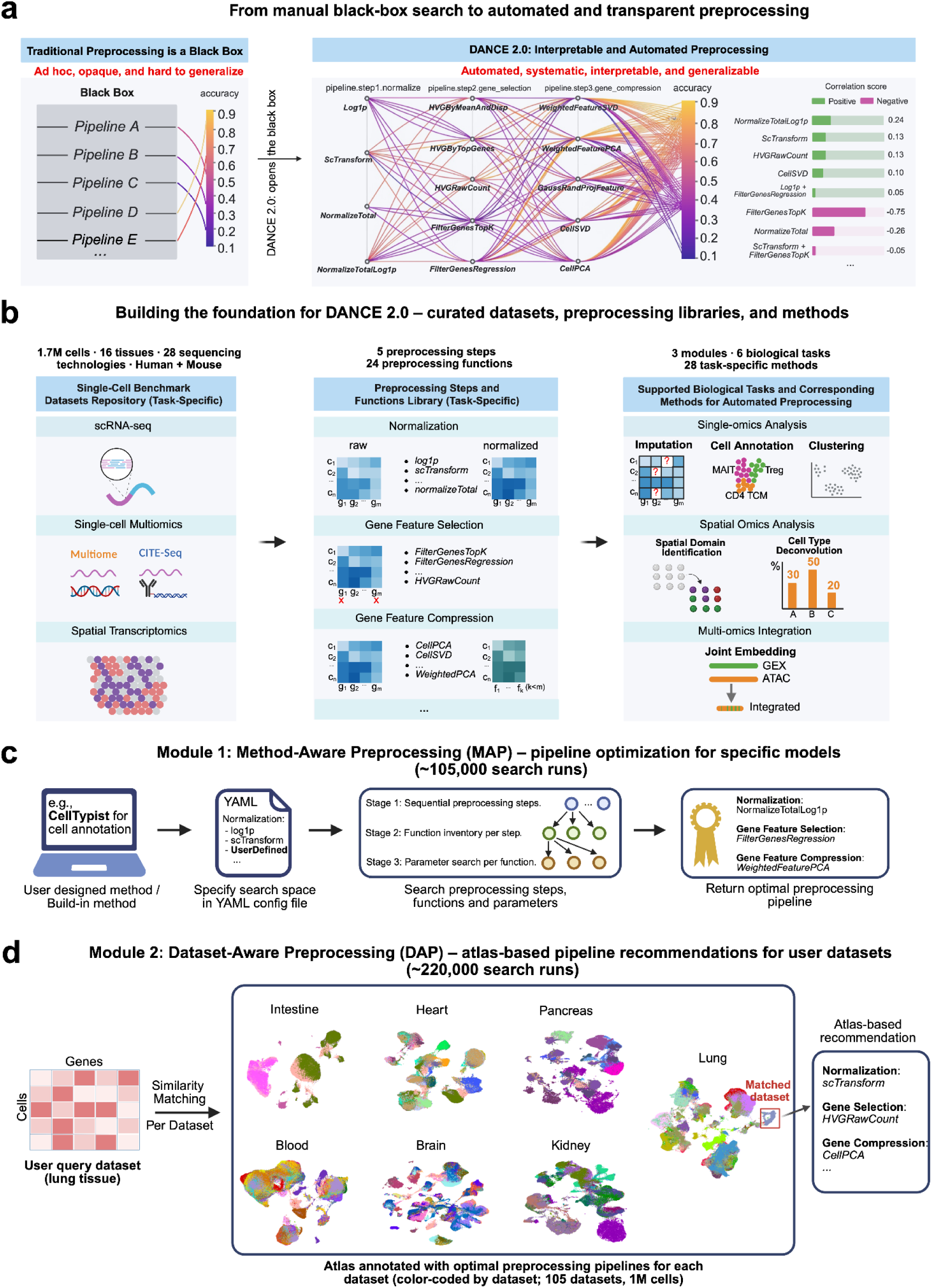
DANCE 2.0 overview: transitioning from opaque manual preprocessing to interpretable and automated workflows for single-cell data and methods. a. **The difference between traditional manual preprocessing and DANCE 2.0: toward interpretable and automated workflows.** Traditional preprocessing is opaque, ad hoc, and difficult to generalize. As shown on the left, it often requires manually trying each pipeline in a trial-and-error fashion without insight into why certain pipelines work better. In contrast, DANCE 2.0 (right) automates the search across preprocessing steps and function candidates, enabling systematic exploration of the pipeline space. The chord diagram illustrates how individual preprocessing functions and their combinations contribute to downstream performance. Beyond automation and visualization, DANCE 2.0 also summarizes the most impactful preprocessing functions by correlating them with model accuracy, offering interpretable insights into preprocessing choices. b. **Construction of the DANCE 2.0 platform: Benchmark datasets, preprocessing library, and supported tasks**. We compile a repository of around 1.7M cells across 16 tissues and 28 sequencing technologies, covering diverse biological tasks (e.g., clustering, cell annotation, spatial deconvolution). The search space for each task consists of up to five preprocessing steps and 24 candidate preprocessing functions. Supported tasks span scRNA-seq, multi-omics, and spatial transcriptomics applications. c. **Method-Aware Preprocessing (MAP): Optimizing pipelines for task-specific methods**. MAP enables automated preprocessing tailored to a task-specific method, which can be either a DANCE 2.0 built-in model (e.g., CellTypist for cell annotation) or a custom user-defined model. Users define the search space by specifying preprocessing steps and function candidates in a YAML configuration file, and can optionally register custom functions within the DANCE 2.0 framework. MAP then performs a structured three-stage search: (1) selecting preprocessing steps, (2) identifying candidate functions for each step, and (3) tuning function-specific parameters. Based on this search, MAP identifies optimal preprocessing configurations for the user’s specified method. In total, MAP executes approximately 105,000 search runs across all supported tasks in DANCE 2.0. d. **Dataset-Aware Preprocessing (DAP): Atlas-guided pipeline recommendations for user datasets**. DAP enables preprocessing recommendations by matching a user’s query dataset (e.g., lung tissue) to annotated datasets in a preprocessing atlas using similarity-based retrieval. The atlas is constructed from approximately 220,000 search runs, where each dataset is labeled with its optimal preprocessing pipelines identified through MAP. This enables fast, on-the-fly recommendations of preprocessing configurations. The current version of the atlas in DANCE 2.0 focuses exclusively on cell type annotation. Extending atlas support to additional downstream tasks remains an important direction for future work.

To support automated and interpretable preprocessing, DANCE 2.0 builds on three foundational components (**Fig. 1b**). (1) Benchmark datasets collection: We curate a comprehensive repository by collecting benchmark datasets from state-of-the-art methods for each task and applying rigorous screening criteria based on biological and tissue diversity, sequencing technology diversity, raw data availability, and ground truth annotations (**Methods**). The resulting repository includes over 1.7 million cells spanning 16 tissues, 28 sequencing technologies, and both human and mouse samples, across scRNA-seq, single-cell multiomics (e.g., Multiome, CITE-seq), and spatial transcriptomics. (2) Preprocessing step and function library: DANCE 2.0 compiles 24 preprocessing functions, drawn from methods used by leading approaches across tasks, organized into five sequential steps, for example normalization, gene feature selection, and gene feature compression. (3) Supported biological tasks and methods: The DANCE 2.0 supports six downstream tasks across three major analysis modules – single-omics analysis, spatial omics analysis, and multi-omics integration – using 28 state-of-the-art methods as task-specific baselines for automated preprocessing. It is important to note that while 28 task-specific methods are built into DANCE 2.0, users can also plug in their own designed methods as well as register custom-defined preprocessing functions into DANCE 2.0 to conduct automated preprocessing pipeline search leveraging the foundation and infrastructure of DANCE 2.0. The platform is open-sourced to facilitate easy extension of benchmark datasets and preprocessing functions, enabling community-driven expansion and adaptability. Together, these components form a robust and extensible foundation that supports the two core modules of DANCE 2.0: Method-Aware Preprocessing (MAP) and Data-Aware Preprocessing (DAP).

The Method-Aware Preprocessing (MAP) module (**Fig. 1c**), which enables automated search and recommendation of optimal preprocessing pipelines tailored to specific task methods. MAP takes as input either a built-in or user-defined method (e.g., CellTypist built-in method for cell type annotation) and a YAML configuration file specifying the search space, including preprocessing steps and candidate functions, such as *log1p*, *scTransform*, or any user-defined operations. If no search space is specified, MAP performs a task-specific default full search using all relevant preprocessing steps and functions curated for that task. MAP then performs a structured three-stage search: (1) exploring combinations of preprocessing steps, (2) selecting candidate functions for each preprocessing step, and (3) optimizing parameters within each function (**Methods**). This design enables scalable, systematic, and interpretable pipeline exploration. For each built-in or user-defined task-specific method, MAP returns an empirically optimized preprocessing pipeline. For example, the optimal pipeline for CellTypist may include *NormalizeTotalLog1P* for normalization, *FilterGenesRegression* for gene feature selection, and *WeightedFeaturePCA* for dimensionality reduction. By aligning preprocessing choices with the unique characteristics and assumptions of the target method, MAP improves model performance in a systematic and interpretable manner. In total, MAP conducts over 105,000 preprocessing pipeline search runs across supported tasks, enabling comprehensive exploration of method-specific preprocessing landscapes. These runs also lay the foundation for uncovering task-, method-, and dataset-level patterns, which are presented in subsequent result sections.

The Dataset-Aware Preprocessing (DAP) module (**Fig. 1d**) recommends optimal preprocessing pipelines for newly user-provided datasets based on atlas-level matching. Unlike the Method-Aware Preprocessing (MAP) module, which tailors preprocessing to a specific method, DAP focuses on the characteristics of the input dataset itself. Given a user query dataset (e.g., lung tissue), DAP performs similarity matching against a curated atlas of 105 annotated datasets spanning over 1 million cells across major tissues. Each dataset in the atlas is associated with its own optimal preprocessing pipeline, obtained through MAP. In total, 220,000 search runs are conducted to construct the atlas. When a query is submitted, DAP identifies the most similar atlas dataset and recommends its corresponding preprocessing pipelines. For example, a lung query dataset may be matched to a similar annotated lung tissue dataset in the atlas and be assigned a recommended pipeline consisting of *scTransform* for normalization, *HVGRawCount* for gene selection, and *CellPCA* for gene compression. The atlas used in this work is specifically built for the cell type annotation task. Extending its support to other downstream tasks, such as clustering, spatial analysis, and multi-omics integration, is an important direction for future work. By leveraging the diversity and structure of the annotated atlas, DAP enables fast, interpretable, and data-driven preprocessing recommendations in real time, without requiring users to perform time-consuming searches.

Taken together, DANCE 2.0 opens the black box of preprocessing for single-cell by enabling transparent and automated preprocessing pipeline recommendations, tailored to task-specific methods through MAP and to new user datasets through DAP.

### MAP consistently improves performance across diverse single-cell tasks

The Method-Aware Preprocessing (MAP) module in DANCE 2.0 is to perform a structured three-stage search to identify optimal preprocessing pipelines for task-specific methods (**Fig. 1a**) (**Methods**). The search space begins with **Stage 1**, where MAP explores different sequences of preprocessing steps, each denoted by a labeled module: ***A*** (Normalization), ***B*** (Gene Feature Selection), and ***E*** (Gene Feature Compression). MAP searches over possible step combinations such as *A, B, E, A->B, A->E, B->E,* or *A->B->E*, allowing flexibility in pipeline structure. The order of preprocessing steps is assumed to be fixed (i.e., normalization precedes feature selection, which precedes compression), ensuring consistency with standard single-cell analysis workflows. In **Stage 2**, MAP enumerates all possible candidate functions within each preprocessing step, for example, *A1* (*normalizeTotal*) and *A2* (*scTransform*) for normalization (Step *A*), and *E1* (*CellPCA*) and *E2* (*CellSVD*) for gene feature compression (Step *E*). In practice, Stages 1 and 2 are jointly executed using a grid search strategy across all valid combinations of preprocessing steps and their associated candidate functions. From this initial search, MAP identifies the TOP *k* (defaulting to *k*=3) high-performing function combinations, which are then passed to Stage 3 for fine-grained parameter tuning. For example, combinations such as *A2*, *A2*->*E1*, and *A2*->*B1*->*E1* may be selected based on performance. In **Stage 3**, MAP performs fine-grained parameter tuning for each selected preprocessing function, such as the number of genes in *A2* (*scTransform*) or the number of components in *E1* (*CellPCA*), yielding the final optimized configuration for each method–dataset pair. For each pair, MAP returns two outputs: (1) the TOP *k* (default *k* = 3) high-level preprocessing function combinations using default parameters, derived from the joint Stage 1&2 search (e.g., *A2*, *A2*->*E1*, and *A2*->*B1*->*E1*), and (2) the top fine-grained pipeline configuration obtained from the full three-stage search with parameter optimization.

To evaluate the effectiveness of DANCE 2.0’s MAP module, we assess its impact on downstream performance across five major single-cell analysis tasks, cell type annotation, imputation, clustering, joint embedding, and cell type deconvolution, spanning single-modality, multi-omics, and spatial transcriptomics modules using 14 state-of-the-art methods (**Fig. 2b-f, Supplementary Fig. 1a-f**). For each method, the model architecture and training procedure keep constant, and only the preprocessing pipeline is varied. We compare four settings: (1) the original implementation, as reported in the respective publications; (2) DANCE 1.0 ^5^, a standardized reimplementation using same preprocessing pipelines with original implementation; (3) DANCE 2.0 with a two-stage search (Stages 1&2: function combination without parameter tuning); and (4) DANCE 2.0 with the full three-stage search (Stages 1-3: including function-specific parameter optimization).

**Figure 2:**
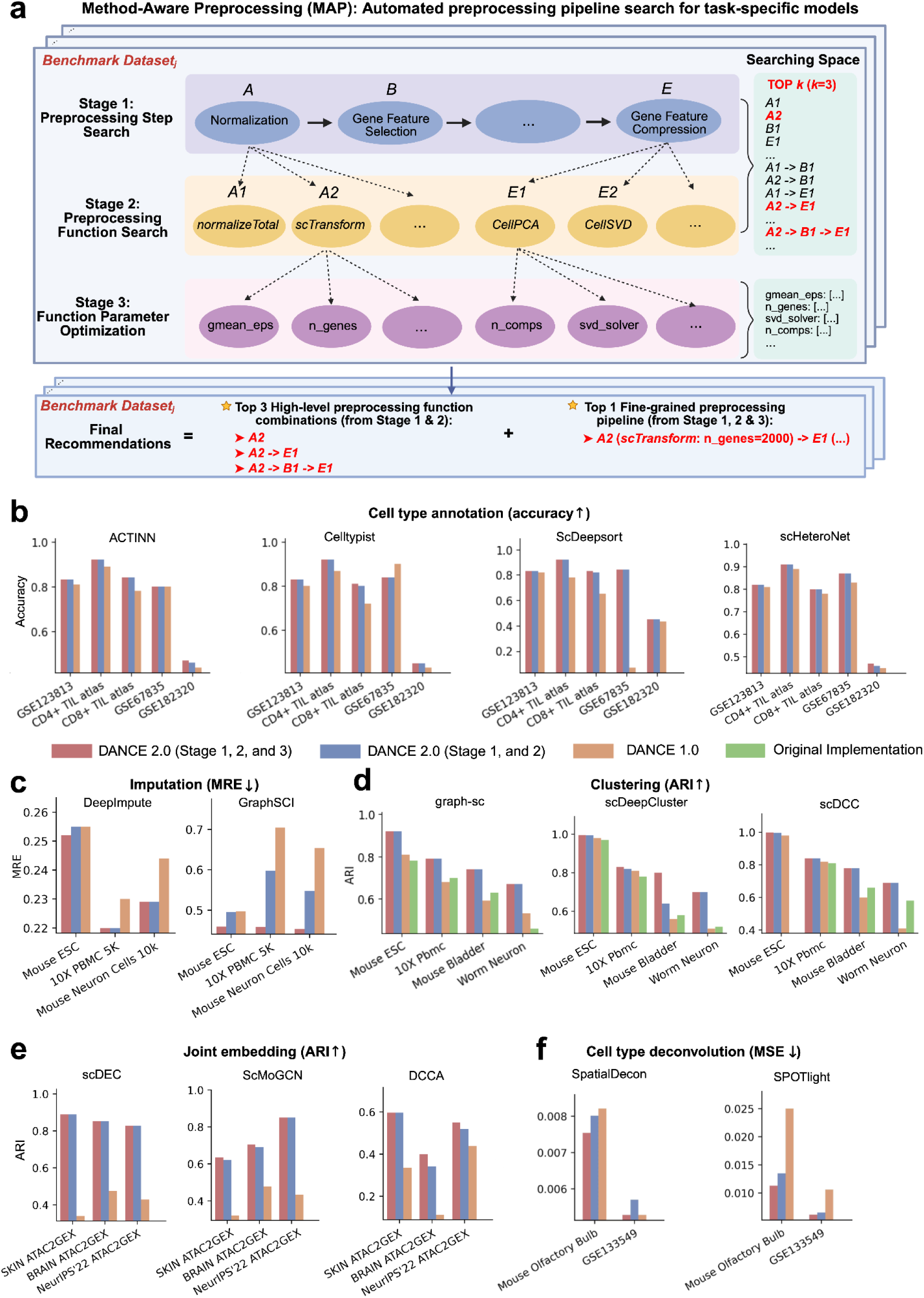
Method-Aware Preprocessing (MAP): automated preprocessing pipeline optimization for task-specific methods and benchmark evaluation across diverse single-cell analysis tasks. a. **The overview of the MAP framework**. DANCE 2.0 performs a structured three-stage search to identify optimal preprocessing pipelines for task-specific methods. Stage 1 explores the sequence of preprocessing steps (e.g., normalization, gene selection, gene compression), Stage 2 searches candidate functions within each step (e.g., *scTransform*, *CellPCA*), and Stage 3 tunes function-specific parameters. Final recommendations consist of two parts: (1) high-level preprocessing function combinations with default parameters, derived from Stages 1 and 2, and (2) fine-grained preprocessing pipelines with tuned parameters, obtained through the full three-stage search. b. **Benchmark comparison of cell type annotation methods under different preprocessing strategies**. Benchmark comparison across four task-specific methods (ACTINN, CellTypist, ScDeepSort, scHeteroNet) under different preprocessing settings. In addition to two DANCE 2.0 configurations including Stages 1&2 (high-level preprocessing function combinations with default parameters) and Stage 1,2&3 (full three-stage search with parameter tuning), we include DANCE 1.0 (orange), a reimplementation of the original method, and the original implementation (green), which reports performance as stated in the original papers. The only difference between DANCE 2.0 and DANCE 1.0 is the preprocessing; the method itself remains unchanged. c. Similar to (b), but for gene imputation. d. Similar to (b), but for clustering. e. Similar to (b), but for joint embedding. f. Similar to (b), but for cell type deconvolution.

In the single-modality module, for cell type annotation, we evaluate four representative methods, ACTINN ^35^, CellTypist ^36^, ScDeepSort ^37^, and scHeteroNet ^12^, across five benchmark datasets (**Fig. 2b**). Overall, preprocessing configurations identified by MAP, including both the full three-stage search (Stages 1-3) and the two-stage variant (Stages 1-2), consistently outperform both DANCE 1.0 and the original method implementations. Notably, DANCE 2.0 achieves substantial accuracy improvements in several settings, such as scHeteroNet on GSE182320 dataset, and scDeepsort on GSE67835 dataset, highlighting the critical impact of preprocessing on downstream performance. For the imputation task, we benchmark DeepImpute ^38^ and GraphSCI ^39^ on three datasets: Mouse ESC, 10X PBMC 5K, and Mouse Neuron Cells 10K (**Fig. 2c**). In all cases, preprocessing pipelines identified by DANCE 2.0, both the high-level combinations from Stages 1-2 and the fully optimized configurations from Stage 1-3, achieve lower mean relative error (MRE) compared to DANCE 1.0 and the original implementations. The most pronounced improvements are observed for GraphSCI and DeepImpute on 10X PBMC 5K and Mouse Neuron Cells 10K datasets. For clustering, we evaluate graph-sc ^40^, scDeepCluster ^41^, and scDCC ^42^ across four datasets: Mouse ESC, 10X PBMC, Mouse Bladder, and Worm Neuron (**Fig. 2d**). Across all methods and datasets, DANCE 2.0 again consistently outperforms baseline settings in terms of Adjusted Rand Index (ARI). Notably, for challenging datasets such as Worm Neuron, DANCE 2.0 preprocessing yields substantially higher ARI scores than both the original implementation and DANCE 1.0, especially for graph-sc, scDeepCluster and scDCC. These results underscore the impact of optimized preprocessing on clustering quality, particularly in noisy or heterogeneous datasets.

In addition to the three tasks in the single-modality module aforementioned, we further assess the generalizability of DANCE 2.0’s MAP in joint embedding from multi-omics module and cell type deconvolution from spatial transcriptomics module. For joint embedding (**Fig. 2e**), we evaluate three representative models, scDEC ^43^, ScMoGCN ^16^, and DCCA ^44^, on three multimodal datasets (SKIN ATAC2GEX, BRAIN ATAC2GEX, and NeurIPS’22 ATAC2GEX). Across all models and datasets, DANCE 2.0 (both full and two-stage optimization) outperforms DANCE 1.0 and the original implementations greatly. For the spatial transcriptomics module, we test SpatialDecon ^45^ and SPOTlight ^46^ on the Mouse Olfactory Bulb and GSE133549 datasets for cell type deconvolution (**Fig. 2f**). DANCE 2.0 achieves substantial reductions in mean squared error (MSE), especially for SPOTlight, where error decreases by more than 50% on the Mouse Olfactory Bulb dataset.

These gains across diverse tasks above are achieved without altering the model architecture or training procedure, only the preprocessing pipeline is optimized. These results demonstrate that MAP in DANCE 2.0 consistently enhances model performance across a wide range of tasks, modalities, and datasets by optimizing only the preprocessing pipeline. This highlights the critical yet often underexplored role of preprocessing in maximizing the effectiveness of single-cell analysis methods.

### MAP preprocessing recommendations generalize across datasets

To assess the robustness of preprocessing pipelines identified by MAP Stages 1 and 2, we design an evaluation framework to quantify their cross-dataset generalizability across multiple benchmark datasets within each task (**Fig. 3a**). For each method-dataset pair, MAP selects the top-*k* pipelines (with *k* = 3, 5, or 10), and we compute the proportion of pipelines that are reused across two or more datasets, serving as a metric for generalizability.

**Figure 3:**
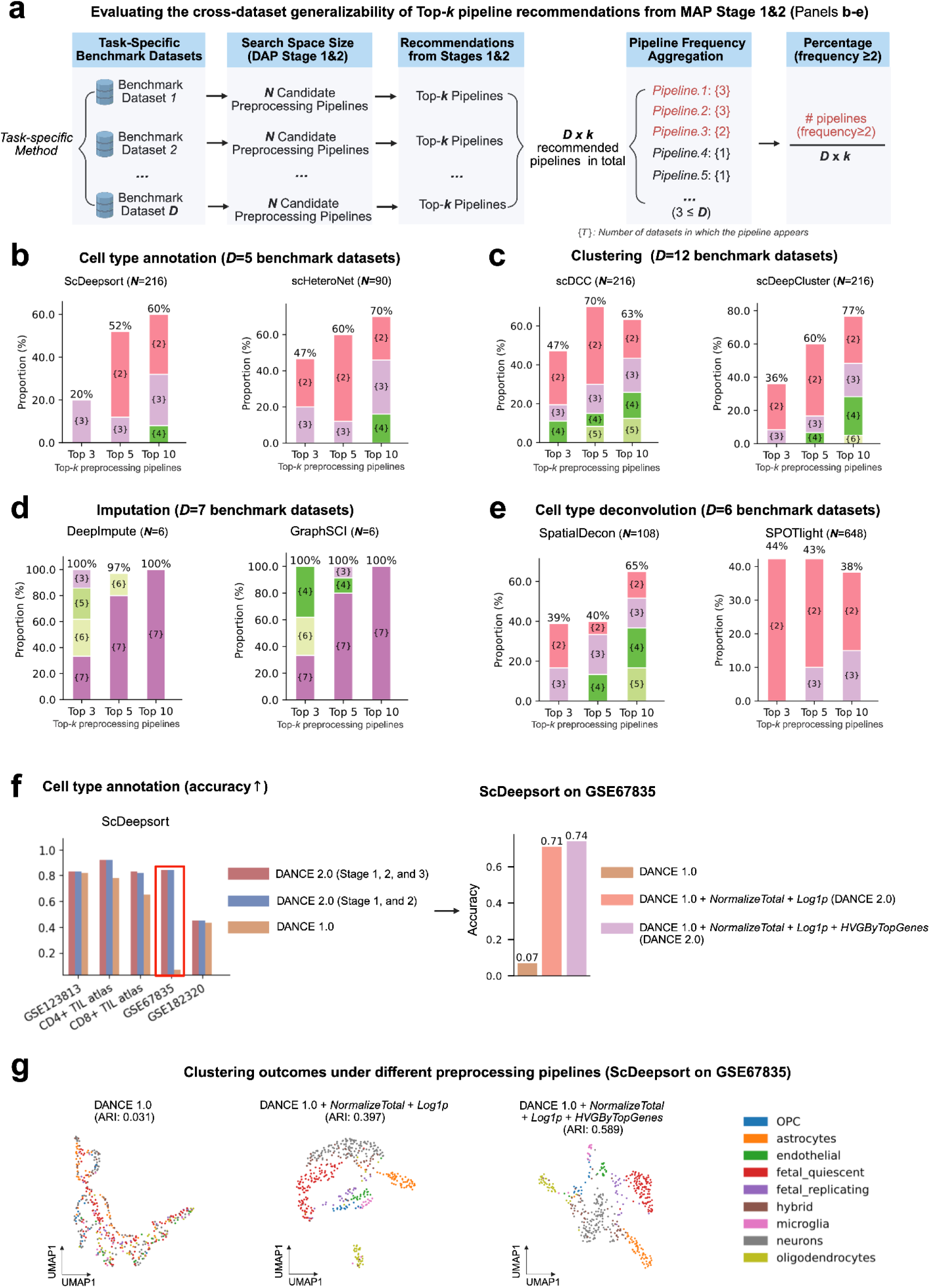
Cross-dataset generalizability and performance dissection of DANCE 2.0 preprocessing pipelines identified by MAP. a. **Evaluation framework for MAP Stage 1&2: Assessing cross-dataset generability of TOP-*k* preprocessing pipelines**. This schematic outlines the evaluation framework for measuring how consistently TOP-*k* preprocessing pipelines, identified through MAP Stage 1&2, are recommended across multiple benchmark datasets for the same task-specific method. For each dataset (***D***), a set of ***N*** candidate pipelines is searched, and the TOP-***k*** performing pipelines are selected. These recommendations are aggregated across all ***D*** datasets, resulting in a total of ***D*** × ***k*** pipelines. Pipeline frequency is computed as the number of datasets in which each pipeline appears. The percentage of pipelines with frequency ≥2 (i.e., reused across at least two datasets) is used to assess generalizability. By default, we set ***k*** = 3, and the number of benchmark datasets per task varies (e.g., D = 5 - 12). b. **Cross-dataset generalizability of TOP-*k* preprocessing pipelines for cell type annotation across benchmark datasets**. Using the evaluation framework defined in (a), we assess the consistency of TOP-*k* preprocessing pipeline recommendations from MAP Stage 1&2 across benchmark datasets for each cell type annotation method. For this task, there are *D* = 5 benchmark datasets. For each dataset, *N* candidate pipelines are searched, *N* = 216 for ScDeepsort ^37^ and *N* = 90 for scHeteroNet ^12^. MAP returns the TOP-*k* performing pipelines (with *k* = 3, 5, or 10) for each method–dataset pair. c. Similar to (b), but for clustering. d. Similar to (b), but for gene imputation. e. Similar to (b), but for cell type deconvolution. f. **Dissecting the performance gains of DANCE 2.0 over DANCE 1.0 through optimal preprocessing from MAP**. **Left**: DANCE 2.0 achieves substantial performance gains over DANCE 1.0, with the most notable improvement observed on the GSE67835 dataset from scDeepsort (highlighted in red). **Right**: To interpret this improvement, we incrementally apply the optimal preprocessing functions identified by DANCE 2.0. Starting from the DANCE 1.0 baseline (accuracy = 0.07), adding *NormalizeTotal* followed by *Log1p* results in a dramatic increase in accuracy to 0.71. Further incorporating *HVGByTopGenes* yields an additional gain, bringing the final accuracy to 0.74. g. **Improved biological structure revealed by optimized preprocessing in ScDeepsort clustering (GSE67835)**. UMAP visualizations show clustering outcomes on the GSE67835 dataset under three preprocessing settings.

Across all tasks, we observe varying degrees of pipeline reuse, with generalizability increasing as the top-k recommendation size grows (**Fig. 3b-e, Supplementary Fig. 2a-e**). For cell type annotation (*D* = 5 datasets), both ScDeepsort ^37^ and scHeteroNet ^12^ show modest reuse among top-3 pipelines (20-47%), which increases substantially at *k* = 10, with 60-70% of pipelines appearing in at least two datasets (**Fig. 3b**). For clustering (*D* = 12 datasets), generalizability is higher overall: scDCC ^42^ and scDeepCluster ^41^ show 36-47% reuse at *k* = 3 and reach 63-77% at *k* = 10 (**Fig. 3c**). For imputation (*D* = 7 datasets), DeepImpute ^38^ and GraphSCI ^39^ exhibit perfect generalizability at all *k* values, with 100% of top-3, top-5, and top-10 pipelines shared across datasets (**Fig. 3d**). For cell type deconvolution (*D* = 6 datasets), SpatialDecon ^45^ and SPOTlight^46^ show moderate but consistent reuse, with 39-44% of top-3 pipelines appearing in multiple datasets (**Fig. 3e**). These results demonstrate that MAP Stage 1&2 reliably identifies preprocessing pipelines that are robust and transferable across datasets, especially as more candidates are considered.

To understand why DANCE 2.0 consistently outperforms DANCE 1.0, we investigate the impact of specific preprocessing choices recommended by MAP. In the cell type annotation task using ScDeepsort ^37^, DANCE 2.0 consistently achieves higher accuracy across all five datasets, with the largest gain observed on GSE67835 (**Fig. 3f, left**). A stepwise breakdown of the improvement on this dataset shows that DANCE 1.0 achieves only 0.07 accuracy. When adding *NormalizeTotal* and *Log1p* (recommended by DANCE 2.0), accuracy jumps to 0.71, and incorporating *HVGByTopGenes* raises it further to 0.74 (**Fig. 3f, right**).

UMAP projections of ScDeepsort’s output on GSE67835 further illustrate the benefit of optimized preprocessing (**Fig. 3g**). DANCE 1.0 produces poorly separated clusters (ARI = 0.031), while the addition of *NormalizeTotal* and *Log1p* improves structure (ARI = 0.397). Incorporating *HVGByTopGenes* results in well-separated, biologically meaningful clusters, achieving ARI = 0.589. These results demonstrate that DANCE 2.0’s performance improvements stem directly from its ability to identify and compose effective, interpretable preprocessing pipelines.

Together, these results highlight that MAP Stage 1&2 not only yields preprocessing pipelines that generalize well across datasets, but also drives substantial and interpretable performance gains. DANCE 2.0’s advantage stems from its ability to systematically identify and compose effective preprocessing steps, ultimately enhancing predictive accuracy and biological resolution across diverse single-cell analysis tasks.

### DAP accurately recommends preprocessing pipelines via atlas-based matching

To enable automated preprocessing for new user datasets, we introduce DAP (Dataset-Aware Preprocessing), a query-to-atlas pipeline recommendation framework (**Fig. 4a**). DAP constructs a reference atlas of 105 scRNA-seq datasets spanning seven main tissue types: blood, brain, heart, lung, intestine, kidney, and pancreas. All datasets selected for the atlas are raw (i.e., unprocessed) and accompanied by cell type labels, ensuring compatibility with supervised cell type annotation evaluation. To ensure both biological and technical diversity, we apply UMAP dimensionality reduction to a range of metadata attributes, including assay type, tissue source, disease status, cell type composition, and sparsity metrics such as total raw counts and non-zero gene counts (**Methods**). This helps to select a representative and diverse subset of datasets for inclusion in the atlas. Each atlas dataset is annotated with four top-performing preprocessing pipelines, identified by MAP Stage 1 & 2 using four cell type annotation methods (ACTINN ^35^, CellTypist ^36^, ScDeepSort ^37^, and singleCellNet ^47^). When a new query dataset is provided, DAP matches it to the most similar datasets in the atlas based on cell-gene expression similarity and recommends the corresponding top pipelines for preprocessing. By default, DAP uses Wasserstein distance ^48^ as the similarity metric for query-to-atlas matching. Additionally, users may select from a suite of eight commonly used distribution- and geometry-based metrics: Wasserstein ^48^, Hausdorff ^49^, Chamfer ^50^, Energy ^51^, Sinkhorn ^52^, Bures ^53^, Spectral ^54^, and Maximum Mean Discrepancy (MMD) ^55^ (**Methods**).

**Figure 4:**
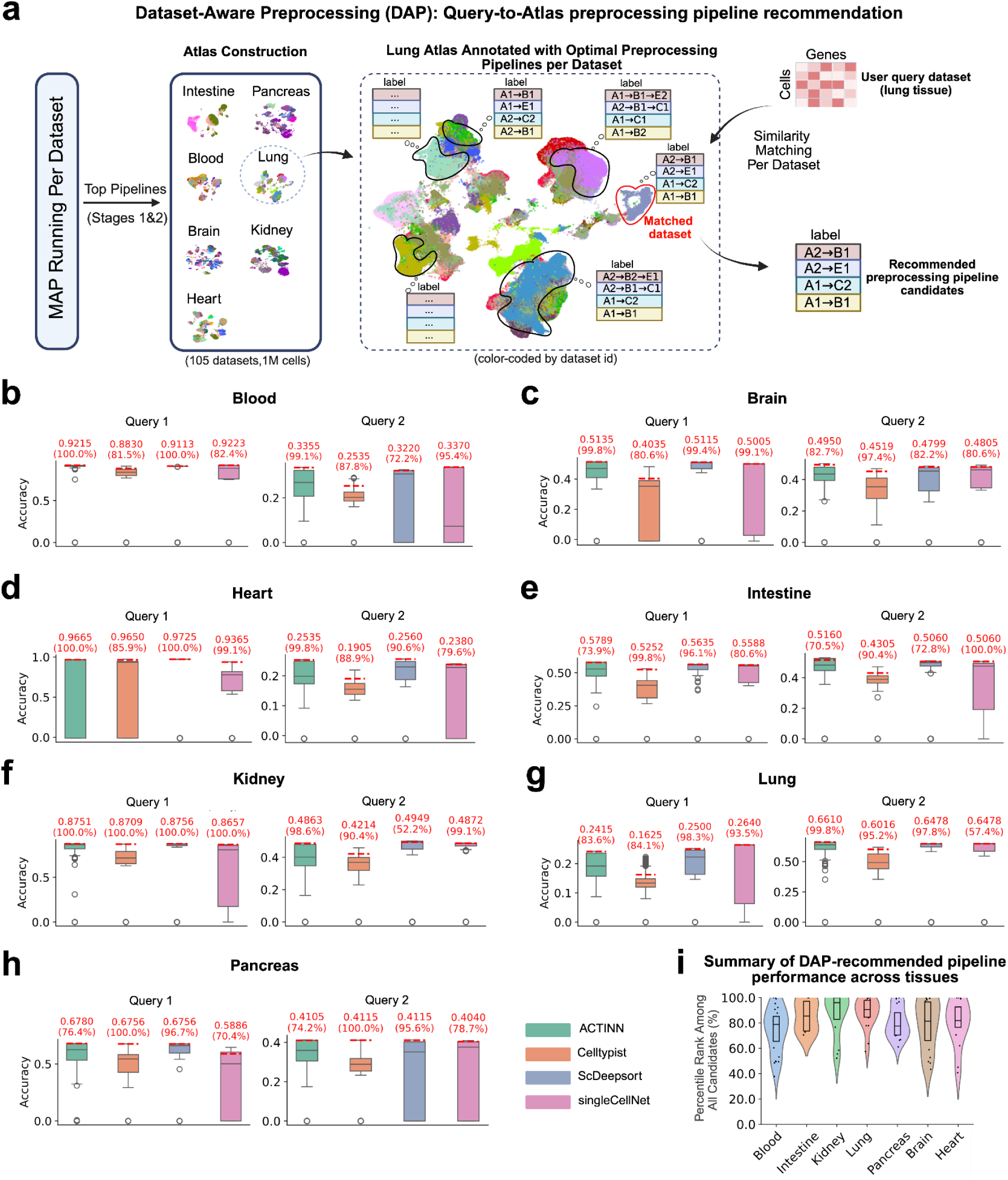
Dataset-Aware Preprocessing (DAP): Query-to-Atlas preprocessing pipeline recommendation and cross-tissue performance evaluation. a. **Overview of the DAP framework**. DAP recommends preprocessing pipelines for a query dataset by matching it to a large reference atlas annotated with optimal pipelines. The reference atlas is constructed using 105 benchmark datasets (1M cells) spanning seven tissues (blood, brain, heart, lung, intestine, pancreas, kidney). Each dataset in the atlas is annotated with its top-performing preprocessing pipelines, identified through MAP Stage 1 & 2 using four cell type annotation methods (ACTINN ^35^, Celltypist ^36^, ScDeepSort ^37^, singleCellNet ^47^). For each method, MAP recommends a single top pipeline, resulting in four annotated pipelines per atlas dataset. When a new query dataset is given, DAP matches it to similar atlas datasets based on cell-gene expression similarity, and retrieves the corresponding top pipelines from matched atlas datasets as recommendations. b. **Effectiveness of DAP-recommended pipelines on blood tissue query datasets**. Each colored box plot represents the distribution of cell type annotation accuracies across all preprocessing pipelines evaluated on the query dataset using a specific method. The red horizontal line indicates the accuracy of the pipeline recommended by DAP, which corresponds to the top-performing pipeline from the most similar dataset in the atlas. The red number above each line shows the accuracy value, and the percentage in parentheses indicates the percentile rank of the recommended pipeline among all candidates, higher values reflect better relative performance. c. Similar to (b), but for brain tissue query datasets. d. Similar to (b), but for heart tissue query datasets. e. Similar to (b), but for intestine tissue query datasets. f. Similar to (b), but for kidney tissue query datasets. g. Similar to (b), but for lung tissue query datasets. h. Similar to (b), but for pancreas tissue query datasets. i. **Percentile rank of DAP-recommended pipelines among all candidate pipelines for each query dataset, grouped by tissue**. Each dot represents a query dataset, and the y-axis indicates the percentile rank of the DAP-recommended pipeline among all candidate pipelines for that query. Higher percentiles indicate stronger performance of the recommended pipeline. Violin plots summarize performance distributions across all queries per tissue, showing that DAP reliably selects high-ranking pipelines across diverse biological contexts.

We assess DAP’s effectiveness using a collection of held-out query datasets across seven tissue types. While each tissue contains multiple held-out queries, we show two representative examples per tissue (**Fig. 4b–h**). Each panel illustrates the performance of four cell type annotation methods across all preprocessing pipelines evaluated on a given query. The red horizontal line indicates the accuracy of the DAP-recommended pipeline, with the corresponding percentile rank reported above each bar. For blood (**Fig. 4b**), DAP-recommended pipelines consistently fall within the 100%-81.5% percentile range for Query 1 and 99.1%-72.2% for Query 2. Similarly high ranks are observed for the brain (**Fig. 4c**), heart (**Fig. 4d**), intestine (**Fig. 4e**), kidney (**Fig. 4f**), lung (**Fig. 4g**), and pancreas (**Fig. 4h**), demonstrating that DAP-recommended pipelines rank among the highest-performing options across all preprocessing pipeline candidates.

To quantify the overall effectiveness of DAP recommendations, we compute the percentile rank of each DAP-recommended pipeline among all candidate pipelines for each query dataset and summarize the results by tissue type (**Fig. 4i**). Each dot represents a single query dataset, and higher percentile values indicate better relative performance of the recommended pipeline. Across all seven tissues, DAP recommendations consistently fall within the top percentile ranges, with median ranks above 80% in most tissues. Notably, kidney, lung, and heart exhibit particularly concentrated high-ranking recommendations, while tissues such as brain and blood show slightly more variability, likely reflecting greater heterogeneity in expression profiles or pipeline sensitivities.

Together, these results demonstrate that DAP robustly generalizes optimal preprocessing strategies to previously unseen query datasets and reliably selects high-performing pipelines across diverse tissues.

### DANCE 2.0 unlocks the black box of preprocessing with task-, method-, and dataset-level interpretability

Most single-cell preprocessing today relies on manual trial-and-error or expert intuition, leading to inconsistent performance and limited reproducibility. This black-box approach lacks transparency, making it difficult to understand why certain pipelines work while others fail. Instead, DANCE 2.0 visualizes all evaluated preprocessing combinations using Sankey diagrams, where each edge represents a unique preprocessing pipeline and is color-coded by downstream clustering performance (Adjusted Rand Index, ARI) (**Fig. 5a**). As shown in two example datasets (Worm Neuron and Mouse Kidney CL2) using the scTAG ^56^ method for clustering, this visualization reveals how specific function choices influence clustering accuracy. For instance, certain normalization approaches like *ColSumNormalize* and gene selection methods like *FilterGenesTopK* often lead to poorer outcomes. This performance landscape enables users to identify and interpret effective preprocessing strategies, offering transparency and diagnostic insight into the impact of each pipeline component.

**Figure 5:**
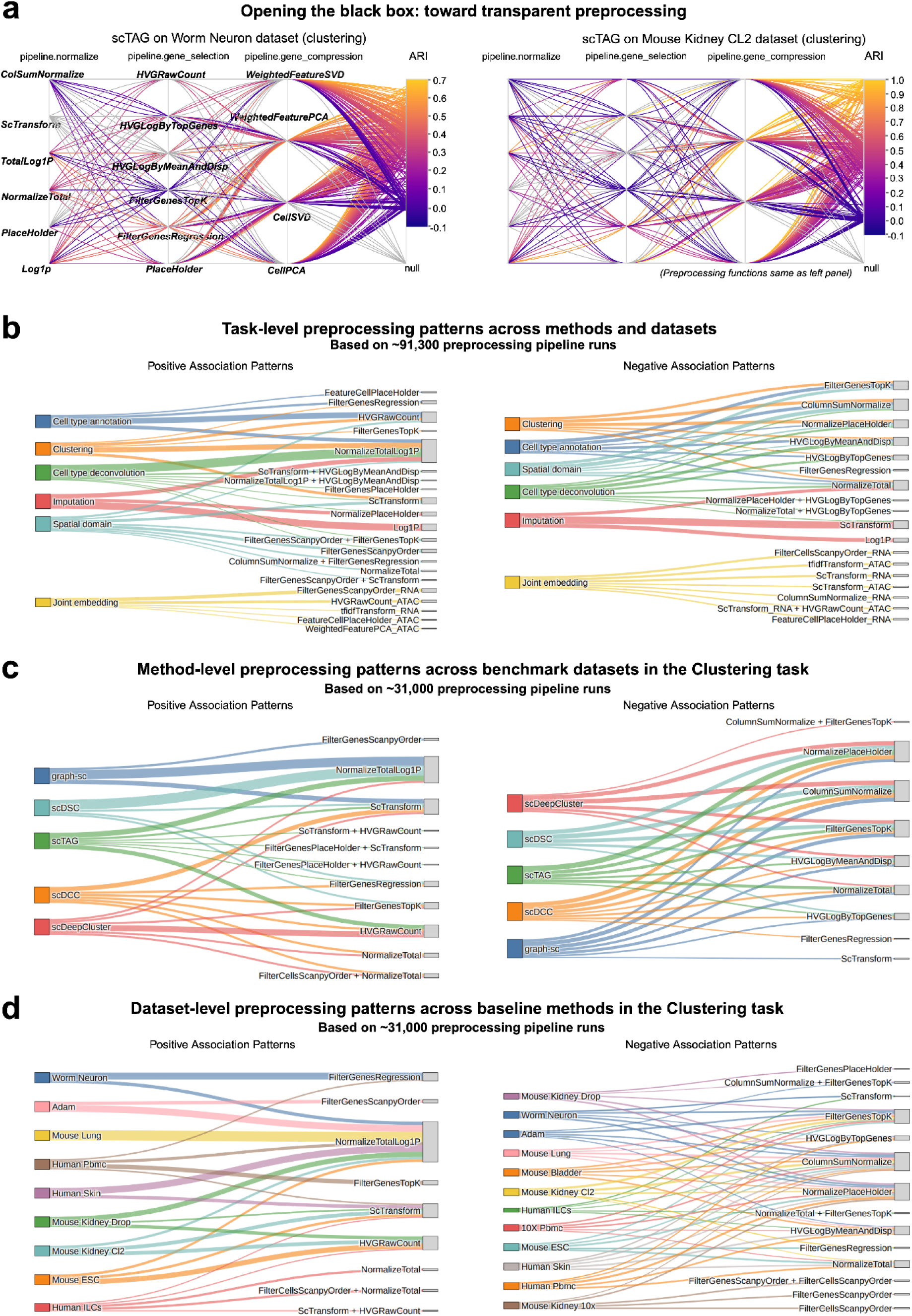
DANCE 2.0 unlocks the black box of preprocessing: dissecting task-, method-, and dataset-level patterns. a. **DANCE 2.0 opens the black box of preprocessing**. Shown are two examples of clustering tasks (Worm Neuron and Mouse Kidney CL2 datasets using scTAG ^56^), where each edge in the Sankey diagram represents a preprocessing pipeline. The color gradient indicates clustering performance (ARI), illustrating how specific combinations of preprocessing functions affect model outcomes. These visualizations provide interpretable links between preprocessing strategies and clustering results. b. **Task-level preprocessing patterns across task-specific methods and benchmark datasets**. Task-level preprocessing associations across methods and datasets, aggregated from 91,300 pipeline runs. Positively (left) and negatively (right) associated preprocessing function combinations are shown for six major task categories (cell type annotation, clustering, imputation, joint embedding, spatial domain identification and cell type deconvolution), illustrating task-specific preprocessing preferences. c. **Method-level preprocessing patterns across benchmark datasets for the clustering task**. Distinct positive and negative associations emerge across five widely used methods (scDCC ^42^, scDeepCluster ^41^, scTAG ^56^, graph-sc ^40^, and scDSC ^57^) under clustering task, revealing clear method-specific preferences and incompatibilities in preprocessing strategies. d. **Dataset-level preprocessing patterns across baseline methods for the clustering task**. Preprocessing function associations at the dataset level reveal unique preferences and incompatibilities across 12 benchmark datasets.

To understand how preprocessing preferences vary across biological tasks, we aggregate task-level patterns from around 91,300 preprocessing pipeline runs (MAP Stages 1&2) spanning six major categories: cell type annotation, clustering, imputation, spatial domain identification, cell type deconvolution, and joint embedding (**Fig. 5b**). For each task, we combine pipeline runs across all supported methods and benchmark datasets in DANCE 2.0, enabling a comprehensive view of task-level preferences. We then identify preprocessing functions that are positively or negatively associated with downstream task performance, based on their consistency across methods and datasets within each task. There are several notable preprocessing patterns across tasks. For example:

**Pattern 1**: *NormalizeTotalLog1P* consistently shows positive associations across all tasks, indicating its general effectiveness as a normalization strategy. In contrast, the negative association patterns for clustering, cell type annotation, spatial domain, and cell type deconvolution exhibit substantial overlap, suggesting shared suboptimal preprocessing choices across these tasks.

**Pattern 2**: *ScTransform* displays task-specific effects, positively associated with spatial domain and clustering tasks, but negatively associated with imputation, highlighting its varying utility depending on the downstream objective.

**Pattern 3**: Normalization methods have a particularly strong influence on imputation performance. Both the positively and negatively associated patterns for imputation are dominated by different normalization strategies, underscoring the task’s sensitivity to this preprocessing step.

To characterize method-specific preprocessing preferences within the task, we take clustering as an example, and aggregate around 31,000 preprocessing pipeline runs across five widely used methods: scDCC ^42^, scDeepCluster ^41^, scTAG ^56^, graph-sc ^40^, and scDSC ^57^ in the task of clustering (**Fig. 5c**). There are several notable preprocessing patterns across tasks. For example:

**Pattern 1**: Methods based on graph neural networks (graph-sc, scTAG, and scDSC) tend to share more positively associated preprocessing pipelines, while autoencoder-based methods (scDCC and scDeepCluster) exhibit a different set of positively associated pipelines. This divergence suggests that model architecture strongly influences the optimal preprocessing strategy, even within the same downstream task.

**Pattern 2**: *NormalizePlaceHolder*, which denotes the absence of any normalization, is negatively associated with all methods. This underscores the critical importance of applying normalization when designing preprocessing pipelines for clustering tasks.

**Pattern 3**: *NormalizeTotalLog1p* and *ScTransform* are consistently positively associated with the most of clustering methods, whereas *ColumnSumNormalize*, *FilterGenesTopK*, and *HVGLogByMeanAndDisp* are consistently negatively associated. This shows that most models benefit from using variance-stabilizing transformations and perform worse with overly simple normalization or strong feature filtering.

To examine how preprocessing preferences vary across datasets, we still take clustering as an example and aggregate results from around 31,000 preprocessing pipeline runs spanning 12 benchmark datasets within the clustering task (**Fig. 5d**). A few notable preprocessing patterns across datasets can be seen below:

**Pattern 1**: *NormalizeTotalLog1p* and *ScTransform* are consistently associated with improved performance across most datasets, while *FilterGenesTopK*, *ColumnSumNormalize*, and *NormalizePlaceHolder* are broadly linked to poor performance.

**Pattern 2**: Human ILCs dataset displays a distinctive set of positively associated preprocessing functions that are not shared with other datasets, underscoring the need for dataset-specific pipeline recommendations - even within the same task. This underscores the critical importance of DAP (Dataset-Aware Preprocessing) in DANCE 2.0, which provides tailored pipeline recommendations based on dataset-specific characteristics, even within the same biological task.

**Pattern 3**: Despite being negatively associated with most datasets, *FilterGenesTopK* shows a positive association with Human Pbmc dataset.

These findings highlight a broader principle: optimal preprocessing choices exhibit both method-specific and dataset-specific dependencies, even within the same task such as clustering. This phenomenon is consistently observed across other major analysis tasks, including cell type annotation, imputation, joint embedding, spatial domain identification, and cell type deconvolution (**Supplementary Fig. 3-7**). These context-dependent patterns underscore the necessity of DANCE 2.0’s intelligent and adaptive design, which supports informed preprocessing decisions tailored to the unique characteristics of both the analytical method and the dataset. By systematically revealing how individual preprocessing function choices affect downstream performance across tasks, methods, and datasets, DANCE 2.0 unlocks the black box of preprocessing. Through the visualization and analysis of over 325,000 evaluated pipeline runs, it exposes both shared and divergent performance patterns, demonstrating the critical need for Method-Aware Preprocessing (MAP) and Dataset-Aware Preprocessing (DAP) strategies to achieve optimal and interpretable single-cell analysis.

## DISCUSSION

DANCE 2.0 reimagines preprocessing in single-cell analysis as a structured, interpretable, and automated process, transforming what has traditionally been a black-box, trial-and-error endeavor into a systematic, data-driven framework. By delivering both method-specific and dataset-specific pipeline recommendations grounded in large-scale empirical evaluation, DANCE 2.0 empowers researchers and practitioners to make informed, reproducible preprocessing decisions tailored to their analytical objectives.

A central innovation of DANCE 2.0 lies in its dual recommendation engine: the Method-Aware Preprocessing (MAP) module and the Dataset-Aware Preprocessing (DAP) module. MAP identifies preprocessing strategies optimized for specific analytical methods, revealing how preprocessing decisions interact with model architectures and training dynamics. In parallel, DAP provides real-time, dataset-specific recommendations by leveraging a curated atlas of over 100 annotated datasets. Together, these modules support both principled benchmarking and practical deployment, with their generalizability and performance gains validated across more than 325,000 pipeline search runs. Critically, DANCE 2.0 goes beyond automation to offer interpretable insights. Our analyses at the task, method, and dataset levels reveal robust and reproducible function-level preferences across thousands of preprocessing pipelines. For instance, transformations such as *NormalizeTotalLog1p* and *ScTransform* consistently lead to improved downstream performance, whereas alternatives like *ColumnSumNormalize* and aggressive gene filtering (e.g., *FilterGenesTopK*) often degrade results. These patterns not only guide effective pipeline design but also inform future method development by clarifying the preprocessing conditions under which specific models perform optimally or fail.

Despite its broad applicability, DANCE 2.0 still has several limitations. First, its function library, while extensive, is limited to commonly used preprocessing operations. Incorporating advanced modules such as batch correction ^58^ may further enhance its utility. Second, although DAP demonstrates strong performance across diverse tissue types and sequencing technologies, its effectiveness ultimately depends on the diversity and representativeness of the reference atlas. Expanding this atlas to include emerging modalities—such as spatial transcriptomics and temporally resolved single-cell data – will be essential to maintaining its relevance and robustness. Third, the current implementation of DAP is specifically designed for the task of cell type annotation. Expanding task-specific atlases to support additional applications – such as genetic imputation, cell type deconvolution, and spatial domain identification – could significantly enhance the breadth and impact of DANCE 2.0.

Looking ahead, DANCE 2.0 provides a flexible foundation for next-generation preprocessing tools. It can be extended to support multi-modal alignment, spatial-temporal single-cell data analysis, and integration with intelligent bio agents ^30–32^ for automated downstream analysis. Importantly, by openly sharing all pipeline search runs – 105,000 from MAP and 220,000 from DAP, DANCE 2.0 contributes a valuable scientific asset for the community. This large-scale, systematic resource enables future meta-analyses, benchmarking studies, and the development of new preprocessing paradigms grounded in empirical evidence.

## ACKNOWLEDGEMENTS

We appreciate D.V.R.P.S (MSU) for assistance with initial data collection, and H.W (MSU) for valuable initial internal discussions during the preparation and execution of this work. We also acknowledge the use of ChatGPT for proofreading the final version manuscript.

## AUTHOR CONTRIBUTIONS

J.D. and Y.W. conceived the study with the goal of automating single-cell preprocessing and led the overall development of the project. Z.X. contributed to method implementation, experiment execution, and data analysis under the guidance of J.D., J.Z., X.Q., J.M., G.Y., and J.T. R.L. implemented YAML configuration support and contributed method plug-in support for the MAP module. J.D. and Y.W. designed and created all figures and drafted the initial version of the manuscript with input from J.M., G.Y., and J.T. J.M., G.Y., and J.T. subsequently refined the manuscript with feedback from all co-authors. All authors read and approved the final version of the manuscript.

## CODE AVAILABILITY

DANCE 2.0 is open-source and freely accessible at https://github.com/OmicsML/dance.

## DATA AVAILABILITY

All datasets used in this study are publicly available, with detailed usage descriptions provided in the **S1 Details About Collected Benchmark Datasets**. The atlas datasets are obtained from the CELLxGENE Census (https://cellxgene.cziscience.com ). For the cell type annotation task, the GSE67835, GSE123813, and GSE182320 (Tissue: Spleen) datasets are downloaded from the Gene Expression Omnibus (GEO). The human CD8+ and CD4+ TIL atlas datasets are available on FigShare at https://figshare.com/articles/dataset/ProjecTILs_human_reference_atlas_of_CD8_tumor-infiltrating_T_cells_version_1/21931875/2 and https://figshare.com/articles/dataset/ProjecTILs_human_reference_atlas_of_CD4_tumor-infiltrating_T_cells_version_1/21981536/1, respectively. For the clustering task, the 10X Pbmc dataset is retrieved from the 10x Genomics website. The Worm Neuron dataset is obtained from the Worm Atlas project (http://atlas.gs.washington.edu/worm-rna/docs/ ), and the Mouse Bladder dataset is hosted on FigShare (https://figshare.com/s/865e694ad06d5857db4b ). The Mouse ESC dataset corresponds to GEO accession GSE65525. The Mouse Lung, Human Skin, and Human Pbmc datasets are downloaded from GEO accession GSE128066, as cited in the scGAC. The Human ILCs dataset is sourced from GEO accession GSE70580, and the Adam dataset from GSE94333. The sc-mixology datasets are retrieved from the GitHub repository at https://github.com/LuyiTian/sc_mixology. For the imputation task, the 10X PBMC 5K and Mouse Neuron Cells 10K datasets are obtained from the 10x Genomics website. Additional datasets include Mouse ESC (GSE65525), Human Breast Cancer (GSE114397), Human Melanoma (GSE99330), and Mouse Visual Cortex (GSE102827), all available via GEO. For the joint embedding task, the BRAIN ATAC2GEX and SKIN ATAC2GEX datasets from SHARE-seq are available under GEO accession GSE140203. The NeurIPS’22 ATAC2GEX dataset is sourced from the Open Problems for Single-Cell Analysis competition on Kaggle (https://www.kaggle.com/competitions/open-problems-multimodal/data). For the spatial domain identification task, the Mouse Posterior Brain, Pancreatic Cancer, and Mouse Brain Anterior datasets were retrieved from the 10x Genomics website. The Human Breast Cancer dataset is described in the original study ^59^. The spatialLIBD Human DLPFC datasets (151507, 151673, and 151676) are available through the spatialLIBD project (https://research.libd.org/spatialLIBD/). For the cell type deconvolution task, the GSE133549 datasets are sourced from GEO. The Mouse Olfactory Bulb dataset is described in the CARD ^60^. The Human Kidney, Liver, and Lung datasets using CosMx SMI are available in SpatialCTD ^2^. All accession numbers and URLs point to publicly available datasets and resources used throughout this study.

## ONLINE METHODS

### Supported tasks for automated preprocessing

DANCE 2.0 builds upon the foundation of DANCE 1.0 ^5^ and extends its capabilities to support a broader set of downstream tasks for automated preprocessing pipeline search. These tasks are organized into three major modules: (i) single-modality module, including cell type annotation, clustering, and gene imputation; (ii) multimodal module, such as joint embedding across data types (e.g., scRNA-seq and protein); and (iii) spatial transcriptomics module, including spatial domain identification and cell type deconvolution. Below, we provide a brief overview of each supported task, along with the corresponding models integrated under each task for automated preprocessing within the DANCE 2.0 platform. For more details on the supported tasks and associated models, please refer to the DANCE 1.0 ^5^.

#### Single-modality module–cell type annotation

Cell type annotation aims to assign cell identity labels by evaluating the similarity between gene expression profiles of unknown cells and those of reference cell types. Given a gene expression matrix, the similarity between a query cell and cells of known types is computed, enabling label inference based on optimal similarity scores. In DANCE 2.0, this task is supported by six models spanning both statistical and deep learning paradigms: scHeteroNet ^12^, ScDeepSort ^37^ (GNN-based), CellTypist ^36^ (logistic regression), SingleCellNet ^47^ (random forest ensemble), ACTINN ^35^ (multilayer perceptron), and SVM ^61^ (support vector machine). Model performance is evaluated using prediction accuracy.

#### Single-modality module–gene imputation

Gene imputation seeks to recover missing or unobserved gene expression values in sparse single-cell RNA-seq data, thereby enhancing data quality for downstream analyses. The task involves predicting likely expression values in dropout-prone regions of the gene-cell matrix. In DANCE 2.0, this task is supported by three models: DeepImpute ^38^ (parallel multi-layer perceptrons for gene-wise prediction), scGNN ^62^ (autoencoder incorporating transcriptional regulatory signals), and GraphSCI ^39^ (dual autoencoders leveraging both cell graphs and expression reconstruction). Model performance is evaluated using Pearson correlation between imputed and ground truth expression profiles.

#### Single-modality module–clustering

Clustering is a fundamental task in single-cell analysis that enables the identification of cell types or subpopulations based on gene expression profiles. The goal is to group cells with similar transcriptional patterns into coherent clusters without requiring prior labels. In DANCE 2.0, this task is supported by five models: graph-sc ^40^ (GNN-based clustering via gene-to-cell graph encoding), scTAG ^56^ (graph autoencoder initialized with KNN graph and TAGCN), scDSC ^57^ (deep structural clustering integrating autoencoder and GNN), scDeepCluster ^41^ (ZINB-based autoencoder model), and scDCC ^42^ (constraint-aware extension of scDeepCluster). Model performance is evaluated using the Adjusted Rand Index (ARI), comparing predicted cluster assignments to ground truth cell types.

#### Multi-modality module–joint embedding

Joint embedding aims to project features from two distinct modalities into a shared low-dimensional latent space, enabling integrated analysis of multi-omics single-cell data. In DANCE 2.0, performance is evaluated using normalized mutual information (NMI) and adjusted Rand index (ARI), based on k-means clustering over the learned embeddings. This task is supported by four deep learning models: scMoGNN ^16^ (GNN-based graph autoencoder with feature fusion), scDEC ^43^ (autoencoder variant leveraging supervised signals), scMVAE ^63^ (multi-omics variational autoencoder), and DCCA ^44^ (dual VAE model maximizing similarity between latent spaces of each modality).

#### Spatial transcriptomics module–spatial domain identification

Spatial domain identification aims to partition tissue sections into biologically meaningful regions based on spatial transcriptomics data, where each spot contains a gene expression profile with associated spatial coordinates. The objective is to cluster these spots into spatial domains that are coherent in gene expression and histological context. In DANCE 2.0, this task is supported by five models: SpaGCN ^64^ (GCN-based model integrating histology and expression), STAGATE ^65^ (graph attention autoencoder using spatial and transcriptomic features), Louvain ^66^ (modularity-based graph clustering), EfNST ^67^ (efficientNet-based model leveraging multi-scale image features to identify fine-grained spatial domains) and stLearn ^68^ (unsupervised method using SME-normalized spatial data for region discovery). Model performance is evaluated using the Adjusted Rand Index (ARI) by comparing predicted clusters to annotated spatial domains.

#### Spatial transcriptomics module–cell type deconvolution

Cell type deconvolution aims to estimate the cellular composition of spatial transcriptomic spots based on aggregated gene expression profiles. As this represents an inverse problem, the task seeks to infer the proportions of distinct cell types contributing to each spot without access to explicit ground truth compositions. In DANCE 2.0, this task is supported by five models: DSTG ^69^ (GCN-based method using mutual nearest neighbors across mixed and reference data), SPOTlight ^46^ (NMF-based decomposition using scRNA-seq references), SpatialDecon ^45^ (non-negative linear regression model assuming log-normal error), STdGCN ^70^ (a graph-based model integrating scRNA-seq and spatial data) and CARD ^60^ (regression-based model incorporating conditional autoregressive priors). Model performance is evaluated using mean squared error (MSE) between the predicted and true cell type proportions.

The tasks and models described above represent the built-in modules currently supported in DANCE 2.0. To ensure flexibility and broad applicability, DANCE 2.0 is designed with a modular architecture that allows users to plugin their own task-specific models and objectives. It is important to note that once integrated, these custom methods can seamlessly leverage the platform’s automated preprocessing pipeline search and evaluation infrastructure to identify the optimal preprocessing pipeline tailored to the user-defined method.

### Benchmark dataset collection across tasks

To ensure that task-specific models with different preprocessing pipelines are evaluated comprehensively, fairly, and reproducibly, we establish a set of rigorous criteria for selecting benchmark datasets across all supported tasks in DANCE 2.0. The benchmark datasets are initially drawn from those used by state-of-the-art methods for each task. To further ensure quality and consistency, we apply additional screening based on the following criteria:

- **Biological and tissue diversity**. Datasets must encompass multiple species (e.g., human and mouse) and a wide range of tissue types, including brain, lung, kidney, and pancreas, to promote broad biological applicability.
- **Sequencing technological diversity**. Selected datasets are required to represent a variety of sequencing platforms, such as 10X Visium, Smart-seq2, MERFISH, and Illumina, reflecting the heterogeneity of real-world experimental protocols.
- **Raw data availability**. All datasets are included in their unprocessed, raw format, excluding any datasets that have undergone any preprocessing like normalization or transformation. This ensures consistency and allows end-to-end evaluation of preprocessing strategies.
- **Ground truth annotations**. Each dataset must provide high-confidence reference labels appropriate to its associated task, enabling quantitative evaluation using task-specific metrics such as accuracy, Adjusted Rand Index (ARI), or mean squared error (MSE).

Based on these criteria, we curated a total of 37 benchmark datasets, spanning three major data modalities:

- 23 scRNA-seq datasets supporting tasks such as cell type annotation, clustering, and gene expression imputation.
- 3 multi-omics datasets (e.g., CITE-seq, SNARE-seq) used for joint embedding of paired modalities.
- 11 spatial transcriptomics datasets supporting spatial domain identification and cell type deconvolution.

All datasets have been fully integrated into the DANCE 2.0 evaluation framework and serve as the foundation for automated preprocessing pipeline benchmarking across the six supported downstream tasks. For more details on the collected benchmark datasets, please refer to **Supplementary 1: Details About Collected Benchmark Datasets**.

### Preprocessing search space design

To support flexible and task-specific optimization, DANCE 2.0 defines preprocessing pipelines as structured sequences of modular operations. We formalize this design as a three-stage hierarchy for the automated preprocessing pipeline search, organized from high to low:

#### Stage 1: Sequential preprocessing steps

For each task, we first compile the set of preprocessing steps commonly employed by state-of-the-art methods specific to that task. These steps are then aggregated into a unified default preprocessing step search space, which serves as the basis for automated pipeline construction across all models associated with the task. For example, the preprocessing steps for cell type annotation include: **Step 1**–gene filtering for quality control, **Step 2**–normalization, **Step 3**–gene feature selection, and **Step 4**–gene feature compression. The exact composition of preprocessing steps may vary slightly by task. It is important to note that the preprocessing steps are applied in a fixed sequential order, and we do not currently consider alternative permutations of preprocessing step order within the search space.

#### Stage 2: Function inventory per step

For each preprocessing step defined in Stage 1, DANCE 2.0 maintains a curated inventory of candidate functions that can be selected during the pipeline search. These functions are collected from widely used single-cell analysis toolkits and pipelines (e.g., Scanpy, Seurat, scvi-tools), as well as from recent state-of-the-art methods. The goal is to provide a diverse and representative pool of preprocessing functions to enable comprehensive preprocessing pipeline search. For example, in the cell type annotation task, the function inventory per preprocessing step includes:

- **Step 1 Gene filtering for quality control**: *FilterGenesPercentile*, *FilterGenesScanpyOrder* and *FilterGenesPlaceHolder*.
- **Step 2 Normalization**: *ColumnSumNormalize*, *ScTransform*, *Log1P*, *NormalizeTotal* and *NormalizePlaceHolder.*
- **Step 3 Gene feature selection**: *HighlyVariableGenesLogarithmizedByMeanAndDisp*, *HighlyVariableGenesRawCount*, *FilterGenesTopK*, *FilterGenesRegression* and *FilterGenesNumberPlaceHolder*.
- **Step 4 Gene feature compression**: *CellPCA*, *CellSVD*, *WeightedFeatureSVD*, *WeightedFeaturePCA, GaussRandProjFeature*, and *FeatureCellPlaceHolder*.

Each task is associated with its own function set, tailored to the nature of the task and the requirements of task specific models. Note that each preprocessing step includes a **PlaceHolder** function, which allows the preprocessing step to be optionally skipped during pipeline search.

#### Stage 3: Hyperparameter search per function

At the most fine-grained level of the search space, DANCE 2.0 supports hyperparameter tuning for each candidate preprocessing function. This enables more nuanced and performance-sensitive optimization beyond the selection of functions alone. The inclusion of hyperparameter search allows DANCE 2.0 to evaluate both coarse and fine-grained preprocessing variations in a unified framework. Each function’s parameter space is registered in the platform’s YAML configuration, enabling full user control over the search space granularity.

This three-stage hierarchical design, spanning step selection, function choice, and hyperparameter tuning, supports comprehensive, extensible, and task-adaptable preprocessing pipeline discovery. For each task, we predefine a three-stage hierarchical search space as the default preprocessing pipeline configuration. Users can further customize and configure the search space at each stage to suit their specific analytical needs. For more details on each preprocessing step and its corresponding set of functions, please refer to **Supplementary 2**: **Details About Preprocessing Search Spaces**.

#### YAML-based configuration

To enable flexible and user-friendly definition of the preprocessing search space, DANCE 2.0 supports configuration via structured YAML file configuration. Each YAML file specifies a task-specific hierarchical search space, including the set of candidate preprocessing steps, the functions permitted at each step, and the associated default or tunable hyperparameters. In the YAML file, users can also configure integration with Weights & Biases (wandb) to track experiment metadata, monitor search progress, visualize pipeline performance, and compare results across configurations in real time. This YAML-based system ensures that users can easily extend, constrain, or modify the preprocessing search space, making DANCE 2.0 adaptable to a wide variety of experimental designs. Please refer to https://github.com/OmicsML/dance for an example of a YAML configuration file.

#### Method-specific customization under each task

In addition to task-level customization, DANCE 2.0 supports method-specific configuration of the preprocessing search space. This design accounts for the fact that different models under the same task may exhibit varying sensitivities or constraints with respect to preprocessing operations. For example, some models require standardized input dimensions, while others may be incompatible with specific normalization strategies or feature compression techniques. These method-specific constraints are implemented through YAML configuration overrides, which are seamlessly incorporated into the pipeline search process. This ensures that each model is evaluated within a preprocessing context that is both functionally compatible and performance-optimized, while maintaining the modular and reusable design of the overall framework. Furthermore, users can register custom preprocessing functions and specify them directly in the YAML file, enabling full extensibility of the search space beyond the built-in function library.

### Method-Aware Preprocessing (MAP): pipeline optimization for specific models

Having previously defined a three-stage hierarchical search space for preprocessing, we now describe how DANCE 2.0 performs automated preprocessing pipeline search and evaluation within this space. In this context, Method-Aware Preprocessing (MAP) refers to identifying the optimal preprocessing pipeline for a given model within a specific task, either one of the built-in methods in DANCE 2.0 or a custom method defined by the user. This section focuses exclusively on model-specific optimization. In contrast, dataset-specific preprocessing recommendations are addressed separately in the next section through Dataset-Aware Preprocessing (DAP).

#### Search algorithm

The search process is grounded in a three-stage preprocessing architecture that organizes the pipeline into conceptually distinct layers. Stage 1 defines the high-level sequence of preprocessing steps, such as normalization, gene feature selection, and gene feature compression, which together determine the overall pipeline structure. Stage 2 maps each of these steps to a set of candidate functions that perform the corresponding transformation, for example, the normalization preprocessing step can be normalize_total or log1p functions. Stage 3 introduces tunable hyperparameters for each function, allowing fine-grained control over function behavior, for example, the base parameter for log1p. This layered design enables flexible, modular, and task-adaptable pipeline configuration.

- **Stage 1&2 preprocessing step and function optimization:** We first enumerate all valid combinations of preprocessing steps from Stage 1 and their corresponding functions from Stage 2. For each preprocessing step (e.g., *A*, *B*, *C*), it identifies all candidate functions (e.g., *A1*, *A2* for preprocessing step *A*; *B1*, *B2* for preprocessing step *B*) and evaluates sequential compositions of preprocessing steps (e.g., *A* -> *B* -> *C*). Within each sequence, all function-level permutations (e.g., *A1* -> *B1* -> *C1*, *A1* -> *B2* -> *C1*) are assessed. Each preprocessing combination is evaluated using a hold-out validation set, where the downstream model is trained on the training set and evaluated on the validation set to compute a task-specific performance metric (e.g., accuracy, ARI, MSE). This evaluation is conducted using a grid search strategy across all combinations. Based on the validation performance, the top K preprocessing combinations (default: K = 3) are selected to proceed to Stage 3 for hyperparameter optimization. For example, if the highest-performing pipelines are *A1* -> *B1* -> *C1*, *A2* -> *B2* -> *C1*, and *A1* -> *B2* -> *C2*, these three combinations will be retained for subsequent fine-tuning of function-specific parameters.
- **Stage 3 preprocessing function parameter optimization**: For each of the top-performing pipelines selected from Stage 1&2, DANCE 2.0 conducts hyperparameter optimization on all constituent functions. This optimization refines the performance of each function by exploring its parameter space. By default, a Bayesian optimization strategy (iterative optimization) is employed, and each pipeline undergoes 10-20 iterations to identify parameter configurations that maximize task-specific performance.

#### Evaluation protocol

To robustly assess the performance of each candidate preprocessing pipeline, DANCE 2.0 employs a hold-out validation strategy. Specifically, the available dataset is split into a training set and a validation set. The downstream model is trained using the training data and subsequently evaluated on the validation set to compute a task-specific metric. Evaluation metrics vary by task and include accuracy (for classification tasks such as cell type annotation), Adjusted Rand Index (ARI) for clustering and spatial domain identification.

#### Visualization and monitoring

To ensure transparency and interpretability of the preprocessing pipeline search and optimization process, DANCE 2.0 provides full integration with the Weights & Biases (Wandb) platform. All pipeline search experiments, including function selection, hyperparameter optimization, and evaluation metrics, are automatically logged and visualized in real time. Users can access comprehensive experiment dashboards via Wandb links. These dashboards provide detailed views of pipeline configurations, performance trends across iterations, model accuracy, and loss curves. In addition to performance tracking, Wandb also monitors system resource utilization (e.g., CPU/GPU usage, memory consumption) and runtime statistics, facilitating in-depth analysis of computational efficiency.

For a given model within a specific task, DANCE 2.0 executes preprocessing pipeline search individually on each benchmark dataset associated with that task. On each dataset, DANCE 2.0 identifies and returns: (i) the top *k* (default *k* =3) preprocessing combinations based on default parameters from Stage 1 & 2, and (ii) the ranked preprocessing combinations with hyperparameters optimized in Stage 3. To identify preprocessing pipelines with strong generalizability, we aggregate the top K default-parameter combinations from Stage 1 & 2 across all benchmark datasets. Combinations that occur most frequently among the top-performing candidates are prioritized, as they demonstrate consistent effectiveness across diverse data scenarios. These frequently selected pipelines are recommended as robust preprocessing configurations suitable for broad and reliable application.

### Dataset-Aware Preprocessing (DAP): atlas-based recommendation for specific datasets

Dataset-Aware Preprocessing (DAP) aims to recommend preprocessing pipelines that are tailored to the characteristics of a new, user-provided dataset. This is achieved through the construction of a preprocessing atlas, which serves as a reference collection of datasets, each annotated with high-performing preprocessing pipelines discovered via Method-Aware Preprocessing (MAP). Unlike MAP, which is computationally intensive and time-consuming, DAP offers an efficient alternative by providing immediate, search-free preprocessing recommendations based on prior knowledge. This enables users to apply well-informed preprocessing pipelines on-the-fly, significantly reducing computational overhead while maintaining robustness and adaptability to new datasets.

Each entry in the atlas is associated with one or more preprocessing pipelines that performed well in MAP-based evaluations, forming the empirical foundation for pipeline recommendation in DAP. While the broader goal of DAP is to develop task-specific atlases for each downstream analysis task (e.g., clustering, imputation, joint embedding), this work focuses exclusively on building up preprocessing atlas for cell type annotation. Extension to additional tasks remains an important direction for future work.

#### Atlas dataset collection

We source human scRNA-seq datasets from the CELLxGENE portal, using the data snapshot released on July 1, 2024. After applying standard quality control procedures, we retain a total of 46.68 million cells spanning 488 datasets, representing seven major human tissue types: blood, brain, heart, intestine, kidney, lung, and pancreas. To ensure both biological and technical diversity, we apply UMAP dimensionality reduction to a range of metadata attributes, including assay type, tissue source, disease status, cell type, and sparsity metrics such as total raw counts and non-zero gene counts. Datasets are stratified by size, large (more than 10,000 cells) and small (fewer than 10,000 cells), and are sampled based on maximal UMAP distance to promote diversity in atlas dataset representation. For tissues with abundant representation (e.g., blood, brain, lung), selective filtering is applied to avoid over-representation, whereas for underrepresented tissues (e.g., intestine, pancreas), all available datasets are included to ensure sufficient coverage. This process yields a final pool of 1,058,783 cells, from which 767,921 cells are used to construct the reference atlas, and 290,862 cells are held out as internal query datasets for validating similarity metrics and evaluating the DAP recommendation framework. These query datasets may later be integrated into the atlas to expand its diversity and coverage. The final reference atlas comprises a balanced distribution of cells across tissues, including 142,068 cells from blood, 124,923 cells from brain, 143,277 cells from heart, 29,164 cells from intestine, 130,420 cells from kidney, 130,661 cells from lung, and 67,408 cells from pancreas. This biologically diverse and well-curated atlas provides a robust foundation for dataset similarity matching and preprocessing pipeline recommendation within the DAP framework.

#### Atlas construction

To build the atlas, we systematically select well-characterized single-cell RNA-seq datasets that span a broad range of tissues, assays, and technical profiles (see “Atlas Dataset Collection”). For each dataset, the full MAP pipeline search is performed, and the top-performing preprocessing pipelines are stored alongside the raw expression matrix and associated metadata. Specifically, for the cell type annotation task, we select the top four optimal preprocessing pipelines per dataset in the atlas, with each pipeline corresponding to one of the four supported annotation models in DANCE 2.0. This ensures method diversity and captures different preprocessing preferences across model architectures. Each atlas entry thus encapsulates a complete record of the preprocessing strategies that yields optimal performance for that dataset and task. The resulting atlas serves as a reference library of empirical preprocessing knowledge, capturing both biological diversity and methodological variation. It is designed to support later retrieval of preprocessing recommendations based on dataset similarity. Importantly, the current version of the atlas is focused exclusively on the cell type annotation task; construction of task-specific atlases for additional downstream applications remains a target for future development.

#### Similarity matching strategy

To enable preprocessing pipeline recommendation based on dataset similarity, we systematically evaluate a suite of candidate similarity metrics to identify the most effective approach for matching new query datasets to entries in the atlas. Specifically, we consider eight metrics commonly used for comparing distributions and geometric structures: Wasserstein, Hausdorff, Chamfer, Energy, Sinkhorn, Bures, Spectral, and Maximum Mean Discrepancy (MMD). Similarity is computed between the cell-by-gene expression matrices of the query dataset and each atlas entry. Notably, these matrices may differ in both the number of cells and the number of genes, reflecting variation in dataset size, gene filtering criteria, and sequencing protocols. The selected similarity metrics are chosen, in part, for their ability to operate on such mismatched, high-dimensional data without requiring explicit alignment or embedding into a shared feature space.

To assess the quality of each similarity metric, we use held-out query datasets for which ground truth cell type annotations are available. For each query, we first perform a full MAP pipeline search to establish a gold standard: the complete set of evaluated preprocessing pipelines and their corresponding downstream classification performance. We then identify the top-matched reference dataset(s) from the atlas using each similarity metric and transfer their recommended preprocessing pipeline(s) to the query. The transferred pipeline’s performance on the query is then compared against the true best-performing pipeline discovered by MAP on the same query dataset.

Based on this evaluation protocol, and informed by theoretical considerations, Wasserstein distance is selected as the default similarity metric in the DAP framework. Importantly, we observe that the specific choice of similarity metric has limited influence on recommendation quality: top-matched atlas datasets show high consistency across different metrics, and the performance gap between transferred and MAP-derived pipelines remains consistently small. This robustness underscores the stability of the atlas-based recommendation strategy, and supports the use of Wasserstein distance as an effective and computationally tractable default for similarity matching in DAP.

### Pattern discovery from preprocessing pipeline experiments

To systematically identify the most influential preprocessing functions and their interactions across large-scale pipeline experiments, we develop a three-phase analytical framework integrating surrogate modeling, importance attribution, and directional correlation analysis. This framework is inspired by hyperparameter importance methods commonly employed in modern AI development platforms, such as Weights & Biases ^71^.

#### Phase I: Surrogate modeling of pipeline performance

Each preprocessing pipeline is represented as a binary feature vector, where each dimension corresponds to a specific function within a defined preprocessing step (e.g., Step 1 Gene filtering for quality control: *FilterGenesScanpyOrder*, *FilterGenesPlaceHolder*, etc; Step 2 Normalization: *Log1P*, *ScTransform*, *ColumnSumNormalize*, etc.). A value of “1” indicates that the function is enabled in the pipeline, while “0” denotes that it is not used. This representation permits multiple functions to be active concurrently across different steps. Using this binary representation feature table, we train a surrogate model, a random forest ^72^, to learn the mapping from preprocessing pipelines to downstream model performance metrics (e.g., accuracy, adjusted rand index, or mean squared error).

#### Phase II: Importance analysis of preprocessing functions

With the surrogate model trained in Phase I, we proceed to quantify the relative importance of preprocessing components using the SHAPIQ framework ^73^. This phase takes as input both the binary feature matrix (indicating which preprocessing functions are active in each pipeline) and the trained random forest surrogate model. SHAPIQ computes unsigned importance scores for individual preprocessing functions (main effects) as well as combinations of functions (interaction effects). These scores identify which functions or function combinations most significantly affect performance, but do not provide information about whether their influence is positive or negative. The most impactful functions and interactions, as ranked by SHAPIQ importance scores, are then selected as input for Phase III. In the subsequent phase, we perform targeted correlation analysis to determine the directionality of their effects on performance.

#### Phase III: Directionality analysis via correlation

To determine whether the most influential preprocessing components positively or negatively affect model performance, we perform targeted correlation analysis using the same inputs as Phase II: the binary feature matrix and the observed performance metrics across pipelines. We compute the Point-Biserial correlation coefficient ^74^ between the performance metric and binary indicators representing the presence of individual functions or function combinations. This approach quantifies whether each component is associated with improved (positive correlation) or diminished (negative correlation) model performance.

This three-phase framework enables systematic interpretation of preprocessing pipeline behavior across large-scale experiments. By combining surrogate modeling with importance attribution and directional correlation analysis, we not only identify which preprocessing functions and combinations are most predictive of model performance, but also characterize whether their influence is beneficial or detrimental. This approach provides interpretable and generalizable insights to guide the design of effective preprocessing strategies in single-cell analysis.

While the aforementioned method is effective for discovering patterns in most tasks, we adopt a specialized approach for imputation with a more constrained search space. Our process begins by selecting the pipelines in the top 40% (representing positive patterns) and the bottom 40% (representing negative patterns) based on their performance. Subsequently, we employ the Apriori algorithm ^75^ on these selected pipelines to identify frequently occurring preprocessing functions and their combinations, which are then defined as the target patterns.

### Implementation details

All analyses are conducted in Python using a suite of well-established frameworks from DANCE 1.0. The core single-cell analysis pipeline is built with Scanpy (v1.10.1), while scikit-learn (v1.3.2) is used for machine learning utilities and evaluation metrics. Deep learning models are implemented in PyTorch (v2.1.1), and Weights & Biases (wandb, v0.16.3) is used for hyperparameter tuning and experiment tracking. Computational experiments are performed on a heterogeneous high-performance computing cluster. The cluster includes nodes equipped with various Intel Xeon CPUs (e.g., Intel Xeon Gold 5318Y, AMD EPYC 7543 32-Core Processor) and NVIDIA GPUs (e.g., NVIDIA GeForce RTX 3090, NVIDIA RTX A5000). Detailed hardware specifications, including CPU and GPU models, core counts, and memory for each experimental run are documented on our public experiment tracking page: https://wandb.ai/xzy11632/dance-dev. To ensure reproducibility, all stochastic components of the pipeline are controlled by fixing random seeds. This included train-valid-test data splits, neural network weight initialization, and dimensionality reduction algorithms such as PCA and UMAP. A consistent seed value of 0 is used across most experiments. Seed values for individual runs are specified in the corresponding scripts, which are available in our public GitHub repository: https://github.com/OmicsML/dance.

## SUPPLEMENTARY NOTES

### S1 Details About Collected Benchmark Datasets

#### Cell type annotation (scRNA-seq dataset)

Cell type annotation is a critical step in single-cell RNA sequencing (scRNA-seq) analysis, involving the assignment of biological identities (e.g., T cell, neuron, epithelial cell) to individual cells or clusters of cells. This process typically relies on the expression patterns of known marker genes, integration with reference atlases, or supervised machine learning models trained on annotated datasets, thereby enabling the biological interpretation of cellular heterogeneity within complex tissues. For the evaluation of single-cell cell type annotation methodologies, we have compiled the following datasets:

- **GSE67835**: The GSE67835 Human Brain dataset provides single-cell RNA sequencing (scRNA-seq) data from a study investigating transcriptome diversity in the human brain ^76^. The study aimed to capture the cellular complexity of both adult and fetal human brain tissue. The data available under this accession specifically includes samples from healthy adult temporal lobe tissue, obtained from epileptic patients undergoing temporal lobectomy. The sequencing was performed using Illumina MiSeq and Illumina NextSeq 500 platforms.The dataset comprises a total of 466 cells and includes 22,088 gene features. The original study successfully classified individual cells into major neuronal, glial, and vascular cell types.
- **CD8+ TIL atlas**: A human dataset comprising 11,021 single-cell transcriptomes of CD8+ tumor-infiltrating T cells (TILs) with 27,421 features ^77^. This reference atlas spans seven cancer types and was generated using semi-supervised STACAS integration.
- **GSE123813**: This human dataset includes 52,121 cells and 23,309 features from patients with basal cell carcinoma, specifically looking at cells pre- and post-anti-PD-1 (cemiplimab) therapy ^78^. The study focused on clonal replacement of tumor-specific T cells and was sequenced using Illumina HiSeq 4000 and NextSeq 500 platforms.
- **CD4+ TIL atlas**: This dataset features 12,631 single-cell transcriptomes from human CD4+ tumor-infiltrating T cells, with 37,330 features ^79^. It is a reference atlas integrated from 20 scRNA-seq datasets covering seven distinct cancer types.
- **GSE182320**: A *Mus musculus* dataset consisting of 35,488 cells and 23,347 features ^80^. This study investigates virus-specific CD4 T cell responses in the context of acute and chronic viral infections (LCMV Armstrong and Clone 13), using Illumina NextSeq 500 and NovaSeq 6000 for sequencing.

#### Clustering (scRNA-seq dataset)

Clustering is an unsupervised computational task designed to group cells with similar characteristics, primarily their gene expression profiles, into distinct clusters. Its primary objective in single-cell studies is to identify and delineate different cell populations or states within a heterogeneous sample, thereby reducing data complexity and revealing intrinsic biological organization. For the evaluation of single-cell cluster methodologies, we have compiled the following datasets:

- **10X Pbmc**: The 10x Genomics PBMC 4K dataset is a publicly available single-cell RNA sequencing (scRNA-seq) resource provided by 10x Genomics ^81^. It features peripheral blood mononuclear cells (PBMCs) from a healthy human donor. The dataset was generated using the 10x Genomics Chromium Controller and their Universal 3’ Gene Expression chemistry. It comprises a total of 4,340 cells and includes data for 16,653 genes. The cells were sequenced on an Illumina HiSeq 4000, and the raw data is available, having been processed with Cell Ranger 2.1.0. This dataset is commonly used for benchmarking and demonstrating single-cell analysis workflows.
- **Worm Neuron**: The Worm Neuron Cell Atlas is a single-cell RNA sequencing (sci-RNA-seq) dataset derived from Caenorhabditis elegans (C. elegans) at the L2 larval stage ^82^. This dataset was generated to create a comprehensive transcriptional map of individual cells in the developing worm. The specific subset referred to as “worm neuron cell” focuses on neuronal cells. It comprises a total of 4,186 cells and includes 13,488 gene features. The sci-RNA-seq technology used allows for high-throughput profiling and is compatible with methanol-fixed cells.
- **Mouse Bladder**: The Mouse Bladder Cell dataset is a single-cell RNA sequencing (scRNA-seq) dataset derived from the Mouse Cell Atlas (MCA) project ^83^. This extensive project aimed to create a comprehensive transcriptomic map of various mouse tissues. The data was generated using Microwell-seq, a high-throughput scRNA-seq method. The specific subset referred to as “mouse bladder cell” contains 2,746 cells isolated from mouse bladder tissue and includes expression data for 20,670 gene features.

- **Mouse ESC**: The Mouse Embryonic Stem Cell dataset, accessible via GEO accession GSE65525, provides single-cell RNA sequencing (scRNA-seq) data from a study ^84^ investigating gene expression variability in mouse embryonic stem cells (mESCs). The study aimed to dissect population heterogeneity and identify distinct subpopulations within mESCs cultured under serum conditions. The scRNA-seq was performed using a modified version of the CEL-Seq protocol. The dataset comprises a total of 2,717 cells and includes measurements for 24,175 gene features.
- **GSE128066**:The GSE128066 dataset originates from a study designed to evaluate the performance of the BD Rhapsody Single-Cell Analysis System for single-cell RNA sequencing (scRNA-seq) across fresh and cryopreserved human and mouse cells ^85^. The dataset includes several distinct biological samples:

◦ **Mouse Lung:** This subset contains 1,756 cells derived from mouse lung tissue, with expression data for 1,000 gene features.
◦ **Human Skin:** This subset, consisting of 4,853 cells, was derived from human skin (epidermal cells and fibroblasts) and includes expression measurements for 1,000 gene features.
◦ **Human Pbmc:** This subset comprises 8,218 peripheral blood mononuclear cells (PBMCs) from human donors, with expression data for 1,000 gene features.
- **Human ILCs** :The GSE70580 dataset provides single-cell RNA sequencing (scRNA-seq) data from a study focusing on human innate lymphoid cells (ILCs) from tonsils and blood ^86^. The study aimed to characterize the heterogeneity and transcriptional landscape of human ILC populations, including ILC1, ILC2, ILC3, and NK cells. The sequencing was performed using the SMART-seq2 protocol. This dataset comprises a total of 648 cells and includes measurements for 64,535 gene features. The research provided a detailed molecular map of human ILCs, revealing distinct transcriptional profiles and potential developmental relationships.
- **Adam**: The Mouse Kidney Cell dataset provides single-cell RNA sequencing (scRNA-seq) data from a study ^87^. The research aimed to characterize the cellular composition and transcriptional heterogeneity of the adult mouse kidney. Using scRNA-seq, the study identified various known kidney cell types as well as novel cellular subtypes. The sequencing was performed on the Illumina HiSeq 2500 platform. The dataset, as specified, comprises 902 cells and includes expression data for 16,468 gene features. This work contributes to a deeper understanding of kidney biology at the single-cell level.
- **sc-mixology**: This benchmark dataset, with raw data available under GEO accession GSE118767 ^88^, involves three human lung adenocarcinoma cell lines (HCC827, H1975, H2228). It includes experiments where single cells from these lines were mixed and processed using CEL-seq2, Drop-seq, and 10X Chromium. The table specifies the following subsets used in our study:

◦ **Mouse Kidney Drop**: 225 cells, 15,127 features.
◦ **Mouse Kidney Cl2**: 225 cells, 15,127 features.
◦ **Mouse Kidney 10x**: 902 cells, 16,468 features.

#### Imputation (scRNA-seq dataset)

In the context of sparse high-dimensional data like single-cell transcriptomics, the imputation task involves estimating and filling in missing or unobserved values, typically zero counts attributed to technical ‘dropout’ or low expression. The primary goal is to mitigate the impact of data sparsity, thereby improving the accuracy and reliability of subsequent biological interpretations and computational analyses. To evaluate single-cell data imputation performance, the following datasets were utilized:

- **10X PBMC 5K**: A human dataset containing 5,247 peripheral blood mononuclear cells (PBMCs) with 33,538 features ^89^, generated using the 10x Genomics platform.
- **Mouse Neuron Cells 10k**: This dataset consists of 11,843 cells from the combined cortex, hippocampus, and subventricular zone of an E18 mouse, with 31,053 features ^90^. It was generated using 10x Genomics v3 chemistry and sequenced on an Illumina NovaSeq.
- **Mouse ESC** : This dataset has been introduced in the clustering section.
- **David’s GSE114397**: The human breast cancer dataset from GEO accession **GSE114397**^91^ was utilized in this study. This dataset profiles the Epithelial-Mesenchymal Transition (EMT) in an HMLE cell line induced by two distinct mechanisms. The single-cell RNA-sequencing data was generated using the inDrop platform and sequenced on an Illumina HiSeq 2500. For our analysis, we separated the data into two subsets based on the induction method:

◦ **The Human Breast TGFb**: This subset corresponds to cells stimulated with TGF-beta (TGFb) to induce EMT. It contains 7,523 cells and 28,910 features.
◦ **The Human Breast Dox**: This subset consists of cells where EMT was induced by Doxycycline (Dox)-inducible overexpression of the Zeb1 transcription factor. It comprises 3,496 cells and 27,059 features.
- **Human Melanoma**: This human melanoma dataset, accessible via GEO accession GSE99330, comprises 8,641 cells with 32,287 features ^92^. The study used single-molecule RNA FISH measurements as a reference to assess the performance of DropSeq and Fluidigm C1 single-cell RNA sequencing platforms on melanoma cells. Sequencing was conducted using the NextSeq 500 platform.
- **Mouse Visual Cortex**: This *Mus musculus* dataset from GEO accession GSE102827 provides data from 8,950 cells and 25,202 features ^93^. It is derived from a single-cell (inDrops) analysis of the primary visual cortex in mice that were dark-reared and then exposed to light for varying durations (0h, 1h, or 4h) to study experience-regulated gene expression. The sequencing was performed on an Illumina NextSeq 500.

#### Joint embedding (multi-omics dataset)

The joint embedding task aims to integrate data from two or more distinct modalities (e.g., transcriptomics and epigenomics) into a single, shared latent space. The primary objective is to learn a unified representation that not only preserves the intrinsic structures within each modality but also captures correspondences across different modalities. A key desired outcome of such an embedding is its ability to facilitate accurate downstream tasks, such as the segregation of known cell types, the quality of which can be quantified by clustering performance metrics like the Adjusted Rand Index (ARI). For the joint encoding and clustering of multi-modal single-cell data, the following datasets were employed:

- **SHARE-seq**: This dataset utilizes the SHARE-seq technology, which simultaneously profiles gene expression (RNA-seq) and chromatin accessibility (ATAC-seq) from the same single cells. The specific subset used in this study comprises 34,774 cells from mouse skin and 3291 cells from mouse brain ^94^.
- **NeurIPS’22 ATAC2GEX**: This dataset is from the “Open Problems - Multimodal Single-Cell Integration” Kaggle competition ^95^. It consists of single-cell multiomics data from mobilized peripheral CD34+ hematopoietic stem and progenitor cells (HSPCs) isolated from four healthy human donors.The chromatin accessibility (ATAC-seq) and gene expression (RNA-seq) modalities from the 10x Chromium Single Cell Multiome ATAC + Gene Expression technology are utilized. The data includes measurements taken at multiple time points.

#### Spatial domain identification (spatial transcriptomics dataset)

The spatial domain identification task aims to computationally delineate and characterize distinct anatomical or functional regions within a tissue sample based on spatial transcriptomics data. These domains are typically defined by unique gene expression signatures, distinct cellular compositions, or specific histological patterns. Identifying such spatial domains is crucial for understanding tissue architecture, cellular heterogeneity in its native context, and spatially regulated biological processes. For the identification and clustering of spatial domains, the following datasets were utilized:

- **Mouse Posterior Brain 10x Visium Data**: The Mouse Posterior Brain dataset is a spatially resolved transcriptomics dataset generated using the 10x Genomics Visium platform ^96^. It originates from mouse posterior brain tissue. The dataset captures gene expression across 3,353 tissue spots, with measurements for 31,053 genes.
- **Human Breast Cancer**: This human dataset contains 167,780 cells and 313 genes, profiled using the Xenium In Situ platform ^59^. The study associated with this data focused on high-resolution mapping of the breast cancer tumor microenvironment in FFPE tissue, integrating single-cell, spatial, and in situ analysis. It allowed for the identification of 17 different cell types and the characterization of molecularly distinct tumor subtypes.

- **Pancreatic Cancer**: This human dataset, generated using the Xenium platform with the Xenium Human Multi-Tissue and Cancer Panel (377 genes, supplemented with 97 additional genes), consists of 190,965 cells from FFPE-preserved pancreatic adenocarcinoma tissue ^97^. It was analyzed using Xenium Onboard Analysis. The dataset enabled the identification of 17 cell clusters, including metaplastic cells and distinct endocrine cell populations.
- **Mouse Brain Anterior** : The Mouse Brain Anterior dataset provides spatially resolved transcriptomics data from an adult C57BL/6 mouse brain, specifically a sagittal anterior section ^98^. This dataset was generated using the 10x Genomics Visium Spatial Gene Expression platform and is available as a public resource from 10x Genomics . It comprises 2,695 spots and includes expression data for 32,285 genes.
- **spatialLIBD Human Dorsolateral Prefrontal Cortex (151507, 151673, and 151676):** These three datasets are part of the foundational spatialLIBD project, providing spatially resolved transcriptomics of the human brain using the 10x Genomics Visium platform ^99^. They originate from postmortem adult human dorsolateral prefrontal cortex (DLPFC) tissue, a region critical for cognitive functions. The original study utilized these datasets to map the molecular architecture of the six-layered human neocortex, creating a vital reference for layer-specific gene expression.

◦ **Sample 151507:** This dataset captures gene expression across **4,226 tissue spots**, with measurements for **33,538 genes**.
◦ **Sample 151673:** As a biological replicate, this dataset contains expression profiles for **33,538 genes** across **3,639 spatial spots**.
◦ **Sample 151676:** This replicate dataset provides a spatially resolved view of **3,460 distinct tissue spots**, also measuring **33,538 genes**.

Together, these datasets constitute a powerful resource for exploring the molecular basis of cortical organization and for benchmarking computational methods in spatial genomics.

#### Cell type deconvolution (spatial transcriptomics dataset)

The cell type deconvolution task aims to computationally estimate the relative proportions of distinct cell types within a heterogeneous biological sample, typically from bulk tissue omics data (e.g., RNA-sequencing). This process generally leverages reference datasets of known cell type-specific expression signatures or profiles to dissect the contributions of each constituent cell type to the overall observed signal. Successful deconvolution provides insights into cellular composition changes in various biological conditions or disease states. For the task of cell type deconvolution, we have assembled the following datasets:

- **GSE133549**: This dataset is derived from human primary pancreatic tissue. It comprises 224 spots. The original study is a part involved various scRNA-seq protocols performed on human, dog, and mouse samples ^46,98^. The data is available as raw counts.
- **Mouse Olfactory Bulb**: This dataset consists of 1,185 spots and 11,176 genes from the mouse olfactory bulb, generated using the 10X Visium platform ^60^. It is a synthetically generated dataset for deconvolution, using single-cell data as a reference.
- **Human Kidney**: The Human Kidney dataset is a spatial transcriptomics dataset, a specific subset within the larger SpatialCTD benchmark resource ^2^. This benchmark was developed to evaluate cell type deconvolution methods in the context of the human immuno-oncology tumor microenvironment. The Human Kidney dataset was generated using NanoString’s CosMx Spatial Molecular Imager (SMI) technology. This particular dataset comprises 240 spots and includes measurements for 982 genes. The broader SpatialCTD dataset, which includes human kidney samples, aims to provide realistic spatial transcriptomic data for deconvolution tasks.
- **Human Liver**: The Human Liver datasets, specifically “human hcc liver” (hepatocellular carcinoma) and “human normal liver,” are components of the SpatialCTD benchmark resource ^2^. These datasets were generated using NanoString’s CosMx Spatial Molecular Imager (SMI) technology. The “hcc liver” subset comprises 2,709 spots and includes expression data for 1,003 genes. The “normal liver” subset consists of 3,447 spots/cells, also with measurements for 1,003 genes.
- **Human Lung** : The Human Lung dataset is a spatial transcriptomics dataset, part of the larger SpatialCTD benchmark resource ^2^. The Human Lung dataset was generated using NanoString’s CosMx Spatial Molecular Imager (SMI) technology. This specific dataset comprises 600 spots and includes measurements for 983 genes.

**CELLxGENE portal**. All datasets used to construct the Atlas for the DAP module are obtained from the CELLxGENE portal. The CELLxGENE Discover is a large-scale, publicly accessible resource for single-cell transcriptomics data, developed and maintained by the Chan Zuckerberg Initiative. It aggregates and standardizes curated datasets from numerous studies worldwide, providing a rich foundation for cellular research. For the construction of our reference atlas, we utilized the CELLxGENE data snapshot released on July 1, 2024. From this release, we initially sourced 488 human scRNA-seq datasets, encompassing 46.68 million cells. Following rigorous quality control and a sophisticated sampling strategy designed to maximize biological and technical diversity (as detailed in the Methods section), a final reference atlas of 767,921 cells was constructed. This atlas is well-balanced across seven major human tissue types: blood, brain, heart, intestine, kidney, lung, and pancreas, establishing a robust and diverse foundation for our DAP framework.

### S2 Details About Preprocessing Search Spaces

#### Main Preprocessing Steps

- **Initial Gene Filtering (Quality Control) (`filter.gene(QC)`)** This initial stage focuses on quality control-based gene filtering. The objective is to remove genes that do not meet predefined quality criteria, which is crucial for the reliability of downstream analyses. Available methods include:

◦ *`FilterGenesPercentile`*: The *`FilterGenesPercentile`* preprocessing step is employed to remove genes based on their summarized expression characteristics across all cells. A similar percentile-based gene filtering approach is utilized as a preprocessing step in cell type annotation methods like ACTINN ^35^ prior to model training. Our implementation first calculates a specified summary statistic for each gene (e.g., total expression count, variance, coefficient of variation, or relative variance, determined by the `mode` parameter). Genes are then filtered out if their summary statistic falls outside a user-defined percentile range. Specifically, genes with a summary statistic below the *`min_val`* percentile (e.g., the 1st percentile by default, removing lowly expressed genes) or above the *`max_val`* percentile (e.g., the 99th percentile by default, removing extremely highly expressed genes) are discarded. This reimplemented method allows for robust filtering of genes exhibiting outlier expression and facilitates its integration into automated search procedures.
◦ *`FilterGenesScanpyOrder`*: The *`FilterGenesScanpyOrder`* preprocessing function provides a flexible framework for applying a sequence of gene filtering criteria by leveraging Scanpy’s underlying filtering utilities ^20^. These core Scanpy gene filtering operations, such as setting thresholds for minimum counts or cell prevalence, are standard components in numerous single-cell analysis pipelines, including those described for methods like scDSC ^57^ and graph-sc ^40^. *`FilterGenesScanpyOrder`* allows users to specify an *`order`* of such operations, for example, filtering by minimum total counts (*`min_counts`*), minimum number of expressing cells (*`min_cells`*), maximum total counts (*`max_counts`*), and maximum number of expressing cells *(`max_cells`*). Each criterion, implemented via Scanpy, is applied sequentially to the data, ensuring that genes are progressively filtered based on these user-defined thresholds. This ordered application can be critical, as the outcome of one filtering step can influence the summary statistics used in subsequent steps.
◦ *FilterGenesPlaceHolder*
- **Cell Filtering (`filter.cell(QC)`)** This step aims to identify and remove low-quality cells or outliers from the dataset based on specific metrics, ensuring that subsequent analyses are performed on viable and representative cells. Available methods include:

◦ *`FilterCellsScanpyOrder`*: The *`FilterCellsScanpyOrder`* preprocessing function provides a flexible framework for applying a sequence of cell quality control criteria by leveraging Scanpy’s underlying filtering utilities ^20^. These foundational Scanpy cell filtering operations, such as setting thresholds for minimum library size or number of detected genes, are integral to many single-cell analysis workflows, including the preprocessing stages of methods like GraphSCI ^39^ and DeepImpute ^38^. *`FilterCellsScanpyOrder`* allows users to specify an *`order`* of such operations, for example, filtering by minimum total counts (library size, *`min_counts`*), minimum number of detected genes (*`min_genes`*), maximum total counts (*`max_counts`*), and maximum number of detected genes (*`max_genes`*). Each criterion, implemented via Scanpy, is applied sequentially to the data, ensuring that cells are progressively filtered based on these user-defined quality thresholds. This ordered application can be important, as the cell population remaining after one filtering step can influence subsequent QC decisions.
◦ FilterCellsPlaceHolder
- **Normalization (`normalize`)** The purpose of normalization is to adjust raw gene expression counts to account for technical biases, such as differences in sequencing depth or capture efficiency between cells, making expression levels comparable across cells. Available methods include:

◦ *`ColumnSumNormalize`*: This is a custom preprocessing function we implemented for column-wise normalization. The operation is defined as dividing each column in the data matrix by its sum (i.e., `X / sum(X, axis=0)`). We specifically designed this function to serve as a contrast to the *`NormalizeTotal`* method. Our hypothesis is that this approach will yield suboptimal performance because it obscures global shifts in expression between genes. Its inclusion in our study is twofold: first, to empirically validate our hypothesis, and second, to test the efficacy of our proposed mechanism for identifying essential pattern within a preprocessing pipeline.
◦ *`ScTransform`*: Data normalization and variance stabilization were performed using the *`ScTransform`* method ^100^, a widely adopted approach for scRNA-seq data. This method models gene expression using a regularized negative binomial regression to estimate model parameters (gene-specific intercepts, slopes related to sequencing depth, and dispersion parameters). It then uses these parameters to compute Pearson residuals, which represent standardized, variance-stabilized expression values. These residuals are less dependent on sequencing depth and better suited for downstream analyses such as dimensionality reduction and clustering. Key parameters controlling the transformation include *`min_cells`* (minimum cell prevalence for a gene), *`n_genes`* (number of highly variable genes to model, if not all), and *`n_cells`* (number of cells to subsample for model fitting, if not all).
◦ *`Log1P`*: The *`Log1P`* preprocessing step was applied to the gene expression count data, an operation equivalent to *`scanpy.pp.log1p`* ^20^. This transformation computes *X’* = log (*X* + 1) , where *X* represents the original counts and log denotes the natural logarithm (unless a different *`base`* is specified). This log-plus-one transformation is commonly used to stabilize variance, reduce the skewness of count distributions, and make the data more amenable to downstream statistical analyses that assume approximately normal distributions.
◦ *`NormalizeTotal`*: The *`NormalizeTotal`* preprocessing step was applied to account for differences in cellular library sizes (total UMI counts or reads per cell), functioning similarly to *`scanpy.pp.normalize_total`* ^20^. This method normalizes the gene expression counts for each cell by dividing by its total counts and then multiplying by a scale factor, known as the *`target_sum`*. If *`target_sum`* is set to 1e6, this procedure yields Counts Per Million (CPM) values. By default, if *`target_sum`* is not specified, cells are normalized to the median of total counts across all cells. An optional *`max_fraction`* parameter allows for the exclusion of very highly expressed genes from the calculation of the normalization factors, preventing them from disproportionately influencing the normalization of other genes.
◦ *`NormalizeTotalLog1P`*: The *`NormalizeTotalLog1P`* preprocessing function was applied, which combines two sequential and commonly used steps in single-cell analysis, a pipeline also seen in workflows such as those described for SpaGCN ^64^ and CellTypist ^36^. First, gene expression counts were normalized for cellular library size using a total count normalization procedure (*`NormalizeTotal`*), scaling each cell to a common *`target_sum`* (e.g., 1e6 for Counts Per Million, or the median total count by default). Optionally, genes with expression exceeding *`max_fraction`* can be excluded from the size factor calculation. Following this, a logarithm-plus-one transformation (*X’* = log (*X* + 1), typically natural log via the *`base`* parameter) was applied to the normalized counts to stabilize variance and reduce data skewness.
◦ NormalizePlaceHolder
- **Highly Variable Gene (HVG) Selection (`filter.gene`)** This step involves identifying and selecting genes that exhibit high cell-to-cell variation in expression. The goal is to focus on genes that are most likely to be biologically informative and drive differences between cell populations, while reducing noise and dimensionality. Available methods include:

◦ *`HighlyVariableGenesLogarithmizedByMeanAndDisp`*: Highly variable genes (HVGs) were identified using the *`HighlyVariableGenesLogarithmizedByMeanAndDisp`* method, which is based on Scanpy’s *`highly_variable_genes`* function ^20^. This approach operates on logarithmized expression data and identifies genes exhibiting higher-than-expected variance relative to their average expression. Genes are typically binned by their mean expression (using *`n_bins`*), and dispersion (or a related variance measure) is calculated and normalized within each bin. Genes with normalized dispersion values falling within a specified range (defined by *`min_disp`* and *`max_disp`*) and mean expression values within another range (*`min_mean`* and *`max_mean`*) are selected as HVGs. The *`flavor`* parameter (e.g., “seurat” or “cell_ranger”) determines the specific methodology for calculating and normalizing dispersion. Optionally, HVG selection can be performed per batch (using *`batch_key`*) to mitigate batch effects by selecting genes that are consistently variable across batches.
◦ *`HighlyVariableGenesRawCount`*: Highly variable genes (HVGs) were identified directly from the raw count data using the *`HighlyVariableGenesRawCount`* method. This function employs Scanpy’s *`highly_variable_genes`* implementation with the “seurat_v3” *`flavor`*, which is designed for count data and uses a Pearson residuals-based approach or a model of variance stabilization ^20^. Instead of selecting genes based on explicit mean/dispersion cutoffs, this method typically ranks genes by their variability and selects the top *`n_top_genes`* (e.g., default 1000). The `span` parameter influences the LOESS fit when estimating variance. Optionally, HVG selection can be performed per batch (using *`batch_key`*) to identify genes consistently variable across batches, acting as a lightweight batch correction. These selected HVGs are then used for downstream analyses.
◦ *`HighlyVariableGenesLogarithmizedByTopGenes`*: Highly variable genes (HVGs) were identified from logarithmized expression data using the *`HighlyVariableGenesLogarithmizedByTopGenes`* method, which interfaces with Scanpy’s *`highly_variable_genes`* function ^20^. This approach first calculates normalized dispersion scores for genes, typically after binning them by mean expression (using *`n_bins`* and a specified *`flavor`* like “seurat” or “cell_ranger”). Instead of applying explicit dispersion cutoffs, it then ranks genes based on these scores and selects the top *`n_top_genes`* (e.g., default 1000) as HVGs. Optionally, HVG selection can be performed per batch (using *`batch_key`*) to identify genes consistently variable across batches. These selected HVGs are then used for downstream analyses.
◦ *`FilterGenesTopK`*: The *`FilterGenesTopK`* preprocessing function was used to select a specific number of genes (*`num_genes`*) based on their summarized expression characteristics. This type of top-k gene selection based on summary statistics is also employed as a preprocessing step in frameworks like scGNN ^62^ to focus on informative features. Our function first calculates a chosen summary statistic for each gene across all cells, with options including total sum of expression (*`sum`*), variance (*`var`*), coefficient of variation (*`cv`*, default), or relative variance (*`rv`*), as determined by the *`mode`* parameter. Based on these summary statistics, the function then selects either the top *`num_genes`* (if *`top`* is True, default) or the bottom *`num_genes`* (if *`top`* is False). This provides a direct way to retain, for example, the 1000 genes with the highest coefficient of variation.
◦ *FilterGenesRegression*: Highly variable or informative genes were selected using the *FilterGenesRegression* method, which implements several regression-based feature selection strategies. The specific strategy is determined by the `method` parameter. The “*enclasc*“ method, which models the relationship between mean expression, dropout rate, and log-transformed mean expression to score genes, was adapted from the preprocessing workflow described in the EnClaSC paper ^101^. Other available strategies include a “*seurat3*“ approach, which models log-variance as a function of log-mean expression (using a polynomial regression) and selects genes with the highest residuals, and an “*scmap*“ option, which models the relationship between log-dropout rate and log-mean expression. Regardless of the chosen method, the top *`num_genes`* (e.g., default 1000) are retained for downstream analysis. These methods aim to identify genes that are biologically informative by deviating from general trends observed across all genes.
◦ FilterGenesNumberPlaceHolder
- **Cell-level Feature Extraction / Dimensionality Reduction `(feature.cell`)** The objective here is to transform the high-dimensional gene expression data into a lower-dimensional space while retaining the most significant biological variance. This facilitates visualization, clustering, and other downstream analyses. Available methods include:

◦ *`WeightedFeaturePCA`*: Dimensionality reduction was performed using the *`WeightedFeaturePCA`* method, an approach adapted from preprocessing strategies described in both graph-sc ^40^ and scDeepSort ^37^. This two-step approach first computes Principal Component Analysis (PCA) on the transpose of the gene expression matrix (genes x cells), effectively identifying *`n_components`* principal components that represent major axes of variation across genes (gene PCs). Optionally, features can be normalized (using *`feat_norm_mode`* and *`feat_norm_axis`*) prior to this gene PCA. In the second step, an embedding for each cell is generated by taking a weighted sum of these gene PCs. The weights for this sum are derived from each cell’s own normalized gene expression values. The resulting cell embeddings capture cellular heterogeneity projected onto the principal axes of gene co-variation.
◦ *`WeightedFeatureSVD`*: Dimensionality reduction was performed using the *`WeightedFeatureSVD`* method. This approach is analogous to the *`WeightedFeaturePCA`* method described previously, with the primary modification being the use of Truncated Singular Value Decomposition (SVD) instead of Principal Component Analysis (PCA) in the first step. Specifically, Truncated SVD is first applied to the transpose of the gene expression matrix (genes x cells) to identify *`n_components`* gene singular vectors. Optionally, features can be normalized (using *`feat_norm_mode`*) prior to this gene SVD. Subsequently, an embedding for each cell is generated by taking a weighted sum of these gene SVD components, using the cell’s normalized gene expression values as weights.
◦ *`GaussRandProjFeature`*: Dimensionality reduction was performed using the *`GaussRandProjFeature`* method, which employs Gaussian Random Projection by wrapping the *`GaussianRandomProjection`* class from the scikit-learn library ^102^. This technique projects the high-dimensional gene expression data onto a lower-dimensional subspace of *`n_components`* using a randomly generated Gaussian matrix. The *`eps`* parameter relates to the desired level of distortion in pairwise distances, as per the Johnson-Lindenstrauss lemma, though *`n_components`* is often the primary parameter set directly. This method provides a computationally efficient way to reduce dimensionality while approximately preserving the structure of the data.
◦ *`CellPCA`*: Principal Component Analysis (PCA) was performed on the gene expression matrix (cells x genes) using the *`CellPCA`* function, which utilizes the PCA implementation from the scikit-learn library ^102^, to reduce dimensionality. This method identifies *`n_components`* principal components (PCs) that capture the largest axes of variation in the data. The *`svd_solver`* parameter was set to [*your_solver, e.g., “auto“*]. PCA is a standard preprocessing step in numerous single-cell analysis workflows, including those employed by methods such as scTAG ^56^, EfNST ^67^, and STdGCN ^70^. The resulting PC scores for each cell were used as the lower-dimensional representation for subsequent analyses, such as clustering and visualization.
◦ *`CellSVD`*: Truncated Singular Value Decomposition (SVD) was applied to the gene expression matrix (cells x genes) using the *`CellSVD`* function to reduce dimensionality. This approach is analogous to the *`CellPCA`* method, with the primary modification being the use of Truncated SVD ^102^ instead of Principal Component Analysis. It decomposes the data to identify *`n_components`* singular vectors that capture the largest axes of variation. The *`algorithm`* parameter was set to [*your_algorithm, e.g., “randomized“*]. The resulting transformed cell coordinates were used as the lower-dimensional representation for subsequent analyses.
◦ *FeatureCellPlaceHolder*

All functions with the suffix ***’placeholder’*** are essentially placeholder functions, **designed to skip this process.** The above outlines the main preprocessing pipeline. However, this pipeline can be **adjusted by adding or removing steps**, depending on the specific task and algorithm.

Building upon the general preprocessing control framework previously outlined, this section details the task-specific preprocessing pipelines implemented for each analytical method. These pipelines were carefully adapted—through modification or omission of certain steps—from the general framework. Such adaptations were necessitated by the distinct objectives of each task and the varying input data requirements of the corresponding methods, ensuring both successful execution and the integrity of downstream analyses.

#### Task-Specific Preprocessing Adaptations

- **Cell Quality Control (QC) Strategy:** Recognizing that cell quality control (QC) must be applied judiciously to prevent undue influence on algorithmic performance, and drawing upon extensive empirical observations at the algorithmic level, we adopted fixed cell QC protocol for all tasks except clustering. This exception was made because cell QC requirements exhibit substantial variability among different clustering algorithms.
- **Modifications for Data Imputation Tasks:** For tasks involving data imputation, the dimensionality reduction preprocessing step was entirely omitted. Furthermore, gene QC parameters were standardized. These decisions were made because both dimensionality reduction and overly aggressive gene filtering could patently interfere with, or introduce bias into, the imputation process itself.
- **Preprocessing Search Strategy for Joint Encoding Tasks:** Joint encoding tasks necessitate the preprocessing of data from multiple modalities prior to their integration. This inherently creates an exceptionally large search space for optimal preprocessing pipeline combinations (864*864=746496). To navigate this complexity efficiently, we adopted a two-stage strategy. Initially, we focused on the mouse brain tissue dataset from GSE140203. Using a Bayesian optimization approach, we performed 2,000 iterations to search for effective preprocessing pipelines on this dataset. Subsequently, the top 600 or 1,000 performing pipelines identified from this extensive search were then applied to the remaining datasets for the joint encoding task. This strategy substantially reduced the overall computational time required for identifying suitable preprocessing configurations across all datasets

#### Selection of default parameters in different tasks

Given the substantial size of our dataset and the fact that Step 2 (Combinatorial Preprocessing Path Exploration) does not involve hyperparameter optimization, we implemented a pragmatic preprocessing strategy for initial function evaluation. For functions possessing default parameter values, these defaults were directly adopted. For functions lacking default parameters, or where defaults might be overly restrictive for a diverse dataset, we established a broad, inclusive parameter range. For instance, given that the primary objective of cell and gene quality control (QC) is to eliminate undesirable cells and genes, we adopt a more lenient selection strategy for these specific preprocessing methods. Examples include configuring FilterGenesScanpyOrder to remove genes expressed in fewer than 1% of cells, and FilterGenesPercentile to discard genes with expression levels in the bottom 1st percentile. Even during subsequent parameter optimization for these QC functions, we deliberately restrict their search space. This approach aims to minimize the likelihood of QC procedures disproportionately affecting the overall algorithm performance.

### S3 Details About Benchmark Metrics

**Accuracy (ACC)** is a fundamental metric for classification tasks, and in this study, it is employed to evaluate the performance of **cell type annotation**. It quantifies the proportion of correct predictions among the total number of instances evaluated, providing a general measure of a model’s ability to correctly identify class labels. ACC is calculated as the number of correct predictions divided by the total number of predictions, or more specifically for binary/multi-class problems, as 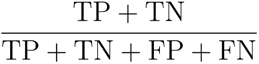, where TP, TN, FP, and FN represent True Positives, True Negatives, False Positives, and False Negatives, respectively. ACC values range from 0 to 1, where 1 signifies perfect accuracy and 0 indicates no correct predictions; thus, higher values are preferable.

**Adjusted Rand Index (ARI)** serves to measure the similarity between two data clusterings. In our work, ARI is specifically applied to evaluate clustering performance across three distinct applications: (1) assessing the quality of clusters derived from **joint embeddings**, (2) evaluating the outcomes of general **clustering tasks**, and (3) measuring the accuracy of **spatial domain identification**. For all these applications, ARI corrects for chance agreement by comparing predicted cluster (or domain) assignments against ground truth labels. It considers all pairs of samples and counts pairs that are assigned in the same or different clusters in both the predicted and true clusterings. The ARI is computed using the formula 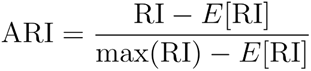, where the “RI” refers to the Rand Index. ARI scores range from -1 to 1: a score of 1 indicates perfect agreement, scores near 0 suggest random agreement, and negative scores imply less agreement than expected by chance. Higher ARI values denote superior quality in these respective clustering and identification tasks.

**Mean Relative Error (MRE)** quantifies the average relative difference between predicted and actual values. For our **imputation tasks**, MRE is used to evaluate the accuracy of the imputed values relative to the ground truth. It is particularly informative when the magnitude of the error relative to the actual value is more critical than the absolute error. MRE is calculated as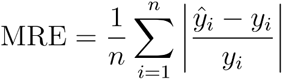 , where `n` is the number of samples, *ŷ_i_* is the predicted (imputed) value for the i-th sample, and *y_i_* is its actual value. Lower MRE values indicate better prediction accuracy, with 0 representing a perfect match. MRE is unitless, but caution is advised when *y_i_* is close to zero.

**Mean Squared Error (MSE)** is a standard metric for evaluating regression-like models, and in this paper, it is applied to assess the performance of **cell type deconvolution**, where the goal is often to estimate proportions or abundances. It measures the average of the squares of the errors—that is, the average squared difference between the estimated values (e.g., cell type proportions) and the actual values. It is calculated using the formula 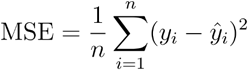 , where `n` is the number of samples (or spots/regions), *ŷ_i_* is the predicted value for the i-th sample, and *y_i_* is its actual value. Lower MSE values indicate a better fit, with an MSE of 0 representing a perfect estimation. By squaring the errors, MSE penalizes larger deviations more heavily.

---

**Supplementary Figure 1:**
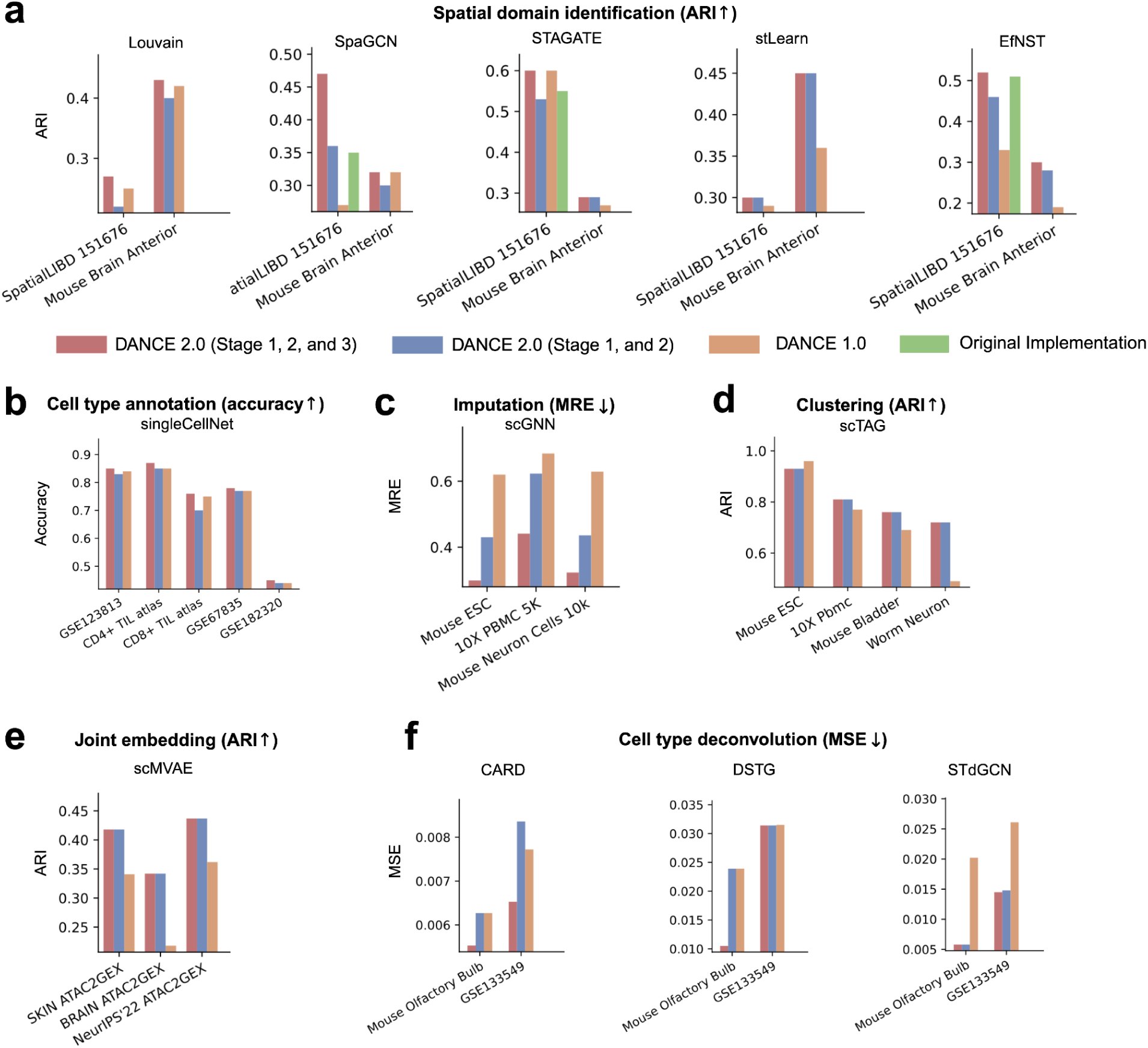
Method-Aware Preprocessing (MAP): automated preprocessing pipeline optimization for task-specific methods and benchmark evaluation across diverse single-cell analysis tasks. a. Benchmark comparison of spatial domain identification methods under different preprocessing strategies. (similar to Fig. 2b). b. Same as in panel a but for cell type annotation. (similar to Fig. 2b). c. Same as in panel a but for imputation. (similar to Fig. 2b). d. Same as in panel a but for clustering. (similar to Fig. 2b). e. Same as in panel a but for joint embedding. (similar to Fig. 2b). f. Same as in panel a but for cell type deconvolution. (similar to Fig. 2b).

**Supplementary Figure 2:**
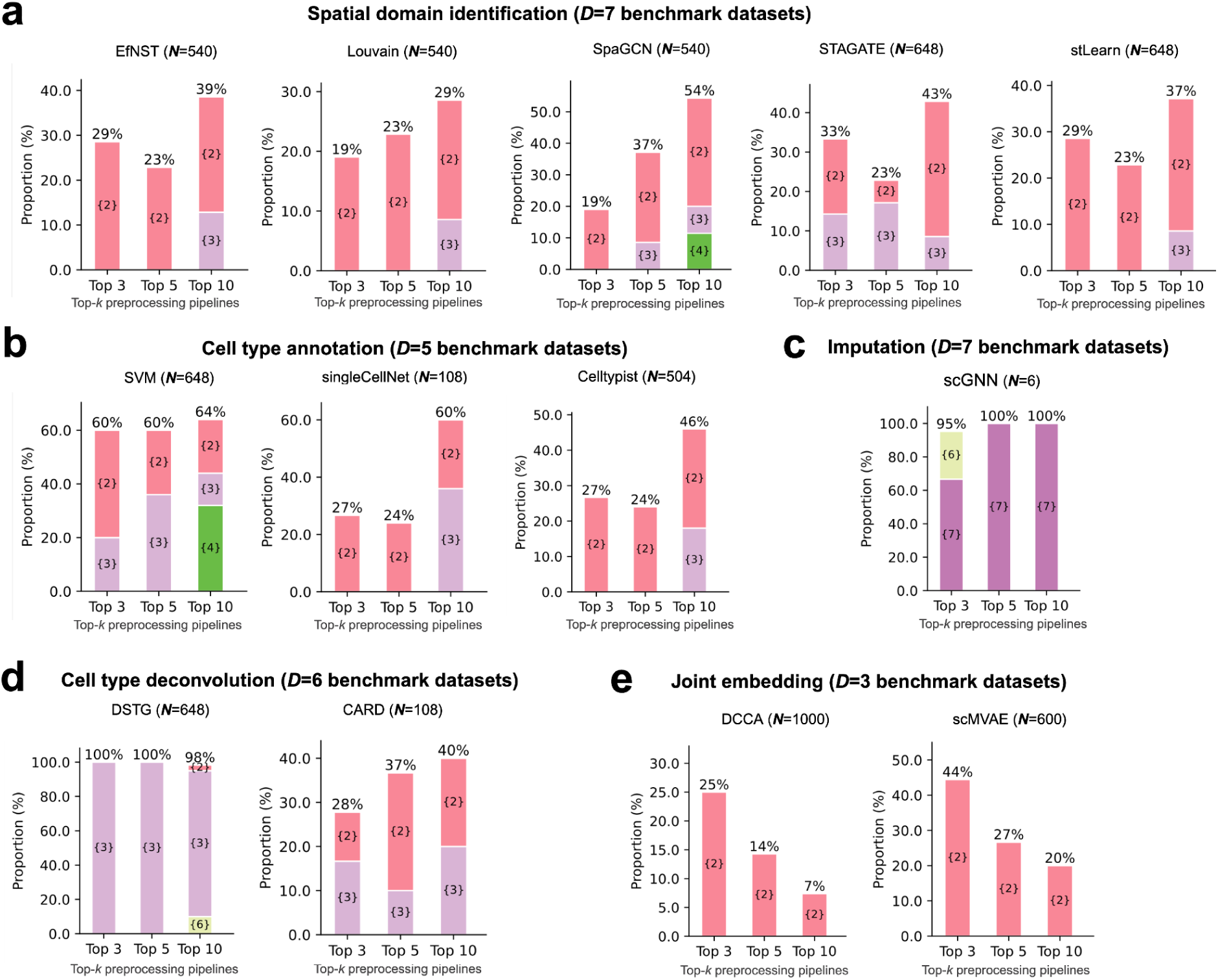
Cross-dataset generalizability of recommended preprocessing pipelines from DANCE 2.0 MAP. a. Cross-dataset generalizability of TOP-*k* preprocessing pipelines for spatial domain identification across benchmark datasets. (similar to Fig. 3b). b. Same as in panel a but for cell type annotation. (similar to Fig. 3b). c. Same as in panel a but for imputation. (similar to Fig. 3b). d. Same as in panel a but for cell type deconvolution. (similar to Fig. 3b). e. Same as in panel a but for joint embedding. (similar to Fig. 3b).

**Supplementary Figure 3:**
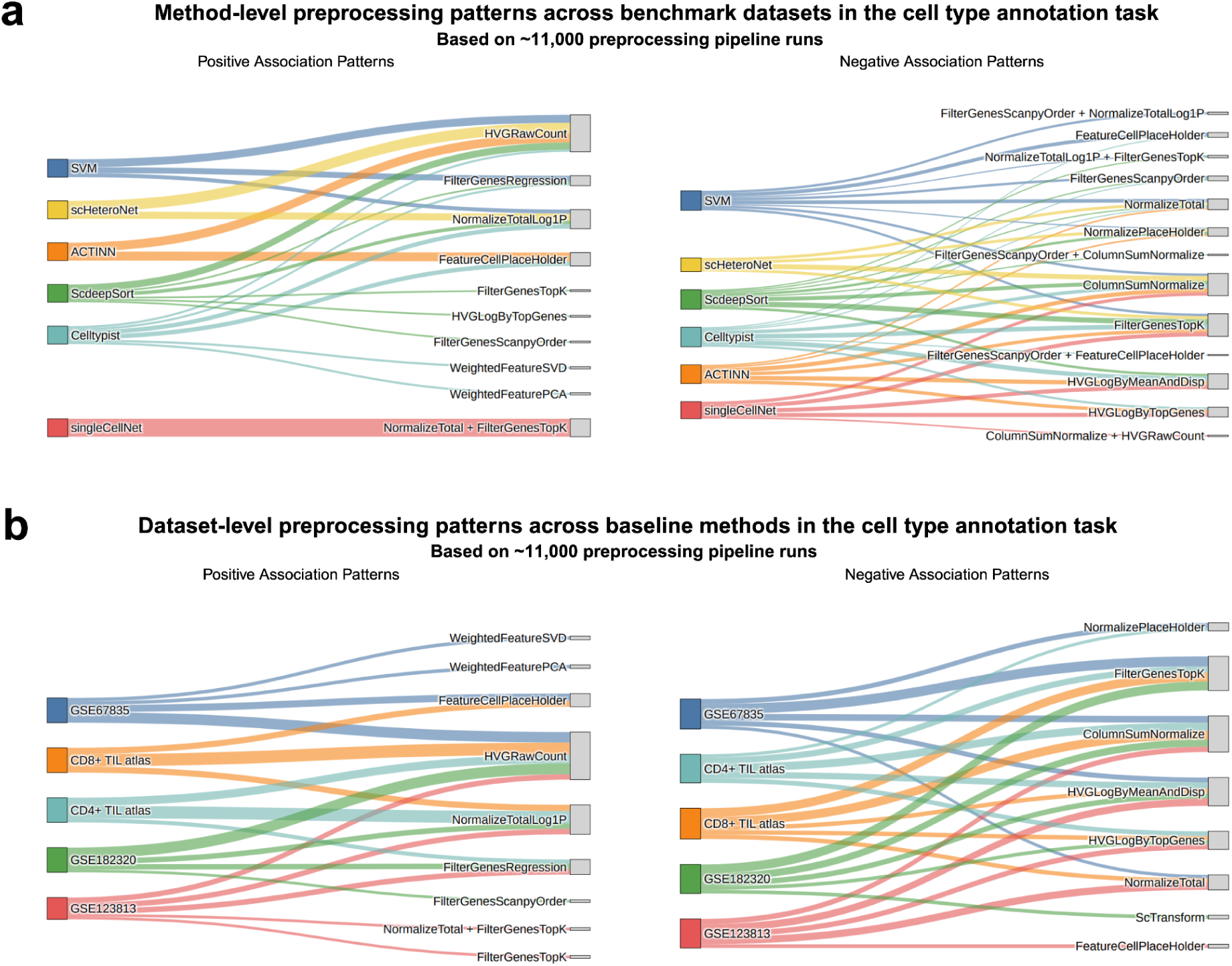
Method- and dataset-level preprocessing patterns identified by DANCE 2.0 in the cell type annotation task. a. Method-level preprocessing patterns across benchmark datasets for the cell type annotation task (similar to Fig. 5c). b. Dataset-level preprocessing patterns across baseline methods for the cell type annotation task (similar to Fig. 5d).

**Supplementary Figure 4:**
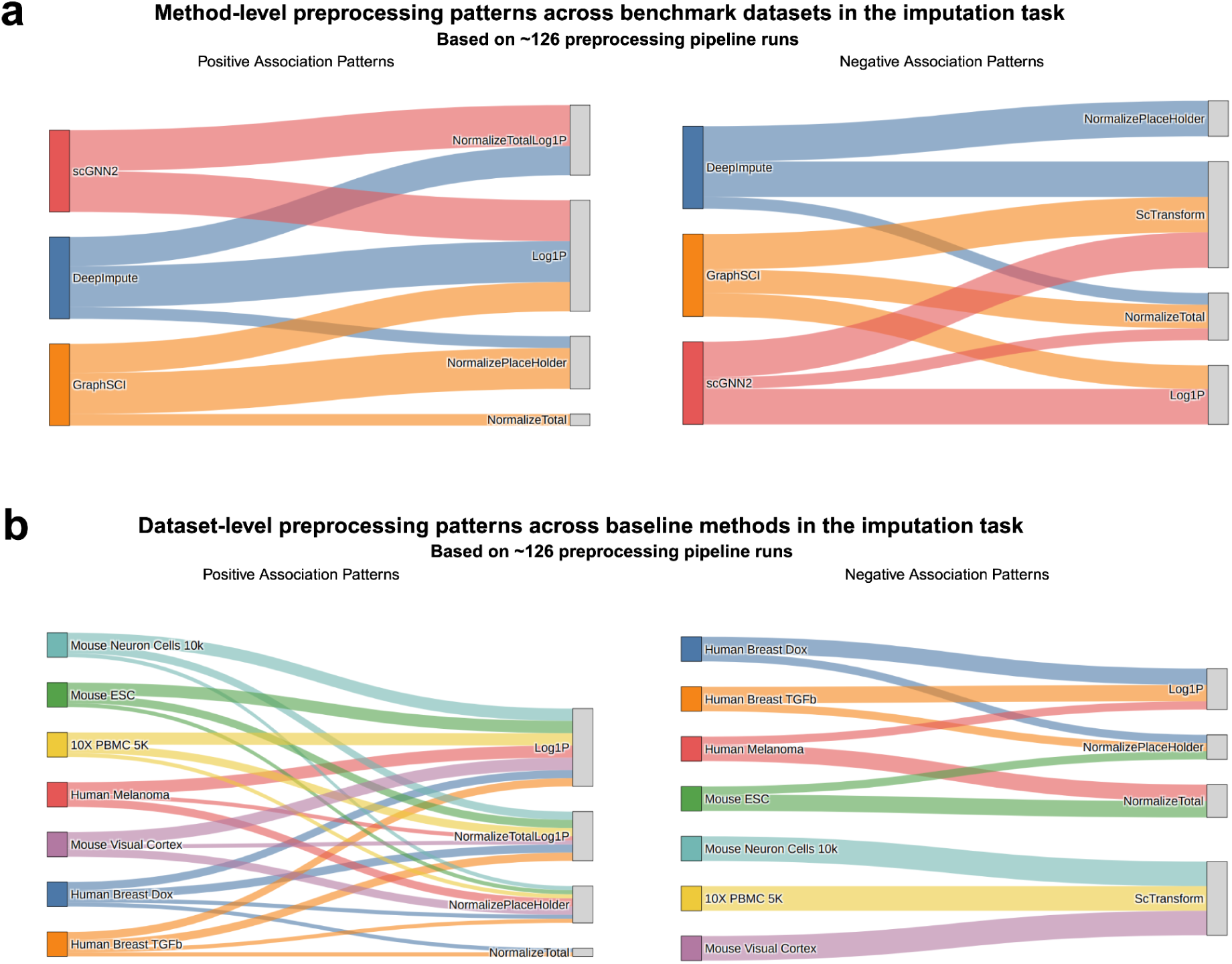
Method- and dataset-level preprocessing patterns identified by DANCE 2.0 in the imputation task. a. Method-level preprocessing patterns across benchmark datasets for the imputation task (similar to Fig. 5c). b. Dataset-level preprocessing patterns across baseline methods for the imputation task (similar to Fig. 5d).

**Supplementary Figure 5:**
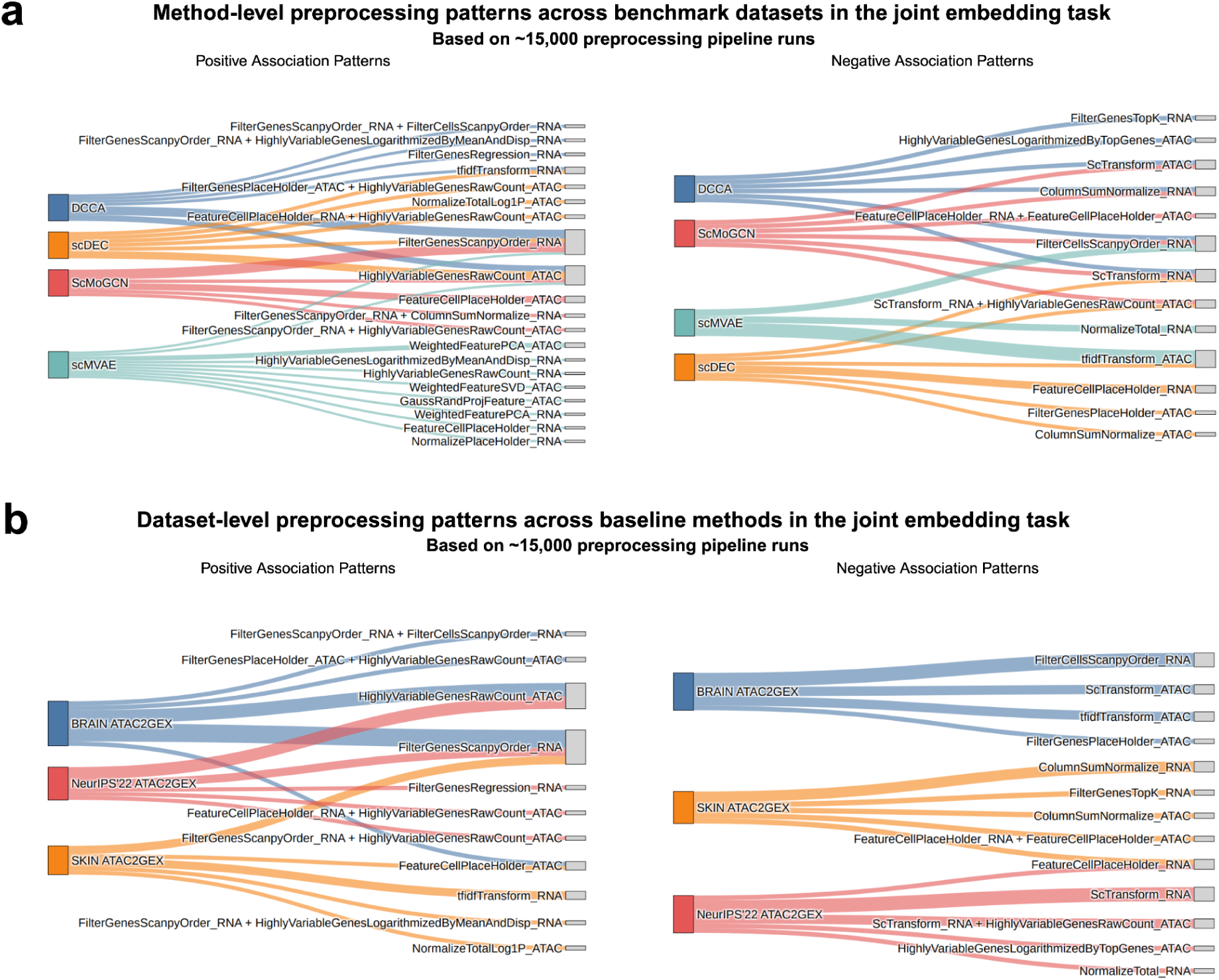
Method- and dataset-level preprocessing patterns identified by DANCE 2.0 in the joint embedding task. a. Method-level preprocessing patterns across benchmark datasets for the joint embedding task (similar to Fig. 5c). b. Dataset-level preprocessing patterns across baseline methods for the joint embedding task (similar to Fig. 5d).

**Supplementary Figure 6:**
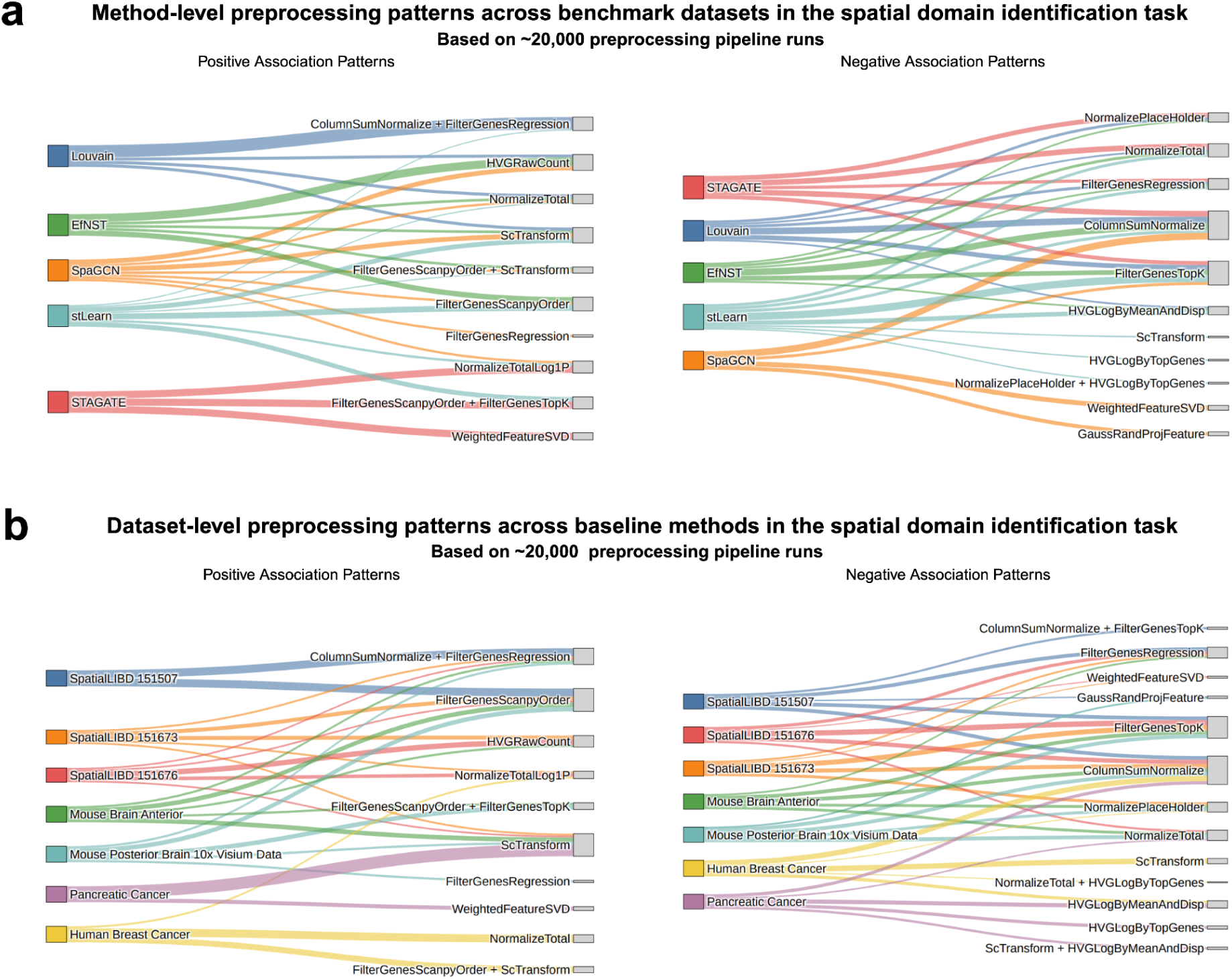
Method- and dataset-level preprocessing patterns identified by DANCE 2.0 in the spatial domain identification task. a. Method-level preprocessing patterns across benchmark datasets for the spatial domain identification task (similar to Fig. 5c). b. Dataset-level preprocessing patterns across baseline methods for the spatial domain identification task (similar to Fig. 5d).

**Supplementary Figure 7:**
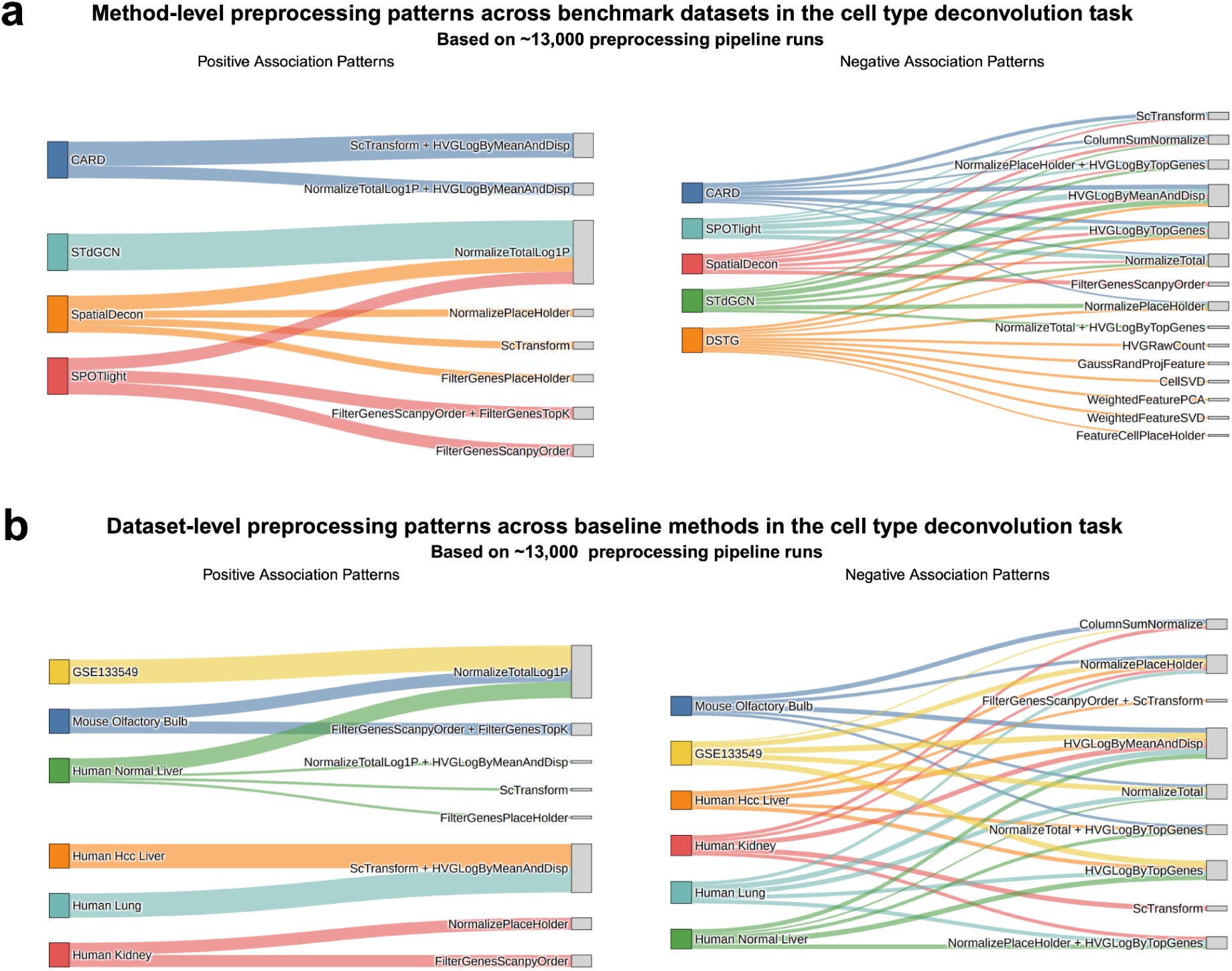
Method- and dataset-level preprocessing patterns identified by DANCE 2.0 in the cell type deconvolution task. a. Method-level preprocessing patterns across benchmark datasets for the cell type deconvolution task (similar to Fig. 5c). b. Dataset-level preprocessing patterns across baseline methods for the cell type deconvolution task (similar to Fig. 5d).

